# Zero-inflated Bayesian factor analysis model with skew-normal priors for modeling microbiome data

**DOI:** 10.64898/2025.12.07.692834

**Authors:** Saurabh Panchasara, Hanna Jankowski, Kevin McGregor

**Affiliations:** Department of Mathematics and Statistics, York University; Department of Statistics, University of Manitoba

**Keywords:** Microbiome data, high dimensionality, additive log-ratio (ALR) transformation, skew normal priors, factor analysis, variational inference

## Abstract

**Background:** Advancements in next-generation sequencing have transformed our understanding of host-microbe interactions, revealing links between microbial composition and chronic conditions such as diabetes, Crohn’s disease, and others. However, the analysis of microbiome data is complex due to its high-dimensionality as well as additional statistical characteristics. One primary objective is to achieve effective dimensionality reduction while simultaneously accounting for the data’s compositional nature and zero inflation. Existing probabilistic models are often based on the assumption that the log-ratio-transformed compositions are normally distributed. This assumption is problematic because it can fail to capture the significant skewness inherent in these transformed compositions.

**Results:** We propose a new model called the Zero-Inflated Factor Analysis Logistic Skew-Normal Multinomial (ZIFA-LSNM) model. ZIFA-LSNM integrates a zero-inflation component to handle excess zeros, employs factor analysis for dimensionality reduction, and, critically, utilizes skew-normal priors on the latent factors to explicitly model data asymmetry. Posterior inference is performed using an efficient variational inference algorithm. Through simulation studies and real data analysis, the ZIFA-LSNM model is shown to have improved performance in parameter recovery and composition estimation compared to its Gaussian-based counterpart.

**Conclusions:** The ZIFA-LSNM model demonstrates that explicitly accounting for skewness in the latent factor structure improves inference in microbiome data. The proposed model, therefore, offers a flexible and scalable framework allowing for the improved analysis of the complex relationships between microbial communities and human health.

## 1. Background

The communities of microorganisms residing on and within the human body, collectively known as the human microbiome (Tyler et al. [2014]), play a pivotal role in regulating host health. These microbial ecosystems influence fundamental physiological processes, from immune system development to metabolism, and have been linked to a wide range of chronic conditions (Ley [2010], Manichanh et al. [2012], Fujimura and Lynch [2015], Lutz et al. [2022]). Advancements in high-throughput sequencing technologies, such as 16S rRNA gene sequencing and shotgun metagenomics, have enabled researchers to understand the composition of these communities in great detail by generating large-scale sample-by-taxon count datasets that are central to modern biological research. However, analyzing these datasets is complex, as they are characterized by a unique combination of statistical properties. They are (i) compositional, (ii) characterized by excess zeros (zero-inflated), and (iii) can exhibit skewness in the log-transformed compositional ratios. Addressing these issues along with the high-dimensionality of the datasets is the major challenge of modern microbiome analysis.

The first challenge in the analysis of microbiome data is its compositional nature. The total number of sequencing reads per sample (often called library size or sequencing depth) is an arbitrary technical artifact, implying that the observed counts provide information only about relative, not absolute, abundances. Failure to account for this proportion-sum constraint can lead to spurious correlations and misleading biological conclusions [Gloor et al., 2017]. The standard approach here is to apply a log-ratio-transformation, which maps the composition from the constrained simplex to an unconstrained real space of one less dimension. These transformations possess the critical properties of scale invariance and sub-compositional coherence, ensuring that analytical results are not affected by observing only a subset of the composition (Van den Boogaart and Tolosana-Delgado [2013]). Early frameworks, such as the Dirichlet-multinomial (DM) and logistic normal multinomial (LNM) models were used as models for the transformed data. However, the DM model cannot adequately capture covariance among taxa due to its limited parameterization (Xia et al. [2013]). To address the correlation structure, Xia et al. [2013] combined a multinomial distribution conditional on latent log-ratio abundances with a Gaussian prior placed on the transformed compositions (Aitchison [1982]).

Next, microbiome data exhibit large numbers of zeros across taxa. Some taxa are truly absent in certain samples (structural zeros), while others go undetected due to limited sequencing depth or experimental biases (sampling zeros) (Lutz et al. [2022]). This high prevalence of zeros creates a major challenge because standard statistical methods, like Poisson or negative binomial models often produce biased results. Several zero-inflated models have been proposed for compositional inference under extreme sparsity (Paulson et al. [2013], Chen and Li [2016], Tang and Chen [2019], Xu et al. [2021], Zeng et al. [2023], Ba et al. [2024]). This challenge is compounded by the “curse of dimensionality,” wherein the number of taxa (*p*) often far exceeds the number of samples (*n*). Consequently, dimensionality-reduction strategies such as probabilistic principal component analysis (PCA) (Tipping and Bishop [1999]), and factor analysis have been frequently incorporated into microbiome models to capture latent community structure.

Recent methodological advances address these challenges individually or in partial combinations. For instance, Zeng et al. [2023] proposed a Zero-Inflated Probabilistic PCA Logistic Normal Multinomial (ZIPPCA-LNM) model, which integrates a low-rank logistic normal framework with a classical probabilistic PCA. To infer the maximum likelihood estimates of latent factors and microbial compositions, they developed an iterative algorithm based on a variational approximation (Ormerod and Wand [2010], Blei et al. [2017], Tran et al. [2021]) within an empirical Bayes framework. More recently, Ba et al. [2024] extended the work of Zeng et al. [2023] to a fully Bayesian setup (ZIPPCA-LPNM). Both models assume Gaussian latent factors.

Despite these advancements, a critical limitation remains in issue (iii) listed above that has not been addressed. As Tu et al. [2024] note : *the log-ratio-transformed compositions of the microbiome data can be highly skewed, especially at a lower taxonomic level. Thus, a Gaussian distribution may not be an ideal prior for the log-ratio-transformed compositions*. Also, *in the presence of skewness in the data, the normal distribution tends to overestimate the number of components by using additional components to capture the skewness (Lin et al. [2007])*. Moreover, Montanari and Viroli [2010] shows that factor models based on the Gaussian assumption can be severely misspecified in the presence of strong asymmetry.

Motivated by this, we introduce the Zero-Inflated Factor Analysis Logistic Skew-Normal Multinomial (ZIFA-LSNM) model, a comprehensive Bayesian framework that unifies the critical statistical challenges inherent in microbiome data. Our approach integrates a zero-inflation component to explicitly account for excessive zeros and utilizes log-ratio-transformation in conjunction with Bayesian factor analysis. In order to capture the asymmetry, commonly observed in additive log-ratio (ALR) transformed data, our model imposes skew-normal priors on the latent factors. This is a key departure from conventional factor analysis, both classical and Bayesian, which typically assumes the latent factors to be standard Gaussian. By introducing skewness into the latent factors themselves, we are able to model a larger spectrum of skewness in the ALR-transformed compositions.

The article is organized as follows. Section 2 formally introduces the ZIFA-LSNM model. To ensure computational efficiency and scalability for high-dimensional datasets, we develop a variational inference algorithm, thereby avoiding the computational expense of Markov chain Monte Carlo (MCMC) methods which often struggle with the complex, multimodal posteriors induced by zero-inflation and skewness. Section 3 presents several simulation studies and demonstrates the model’s application to the real dataset. To motivate and understand the use of skew-normality in our ZIFA-LSNM framework, we conduct a simulation study examining how skewness in the latent space propagates through the full compositional pipeline to the observed log-ratio-transformed counts. As a result, we have show in model simulation results (Section 3.2) and in real data analysis (Section 3.3) that employing a skew-normal distribution improves model fit and yields more interpretable latent structures. This flexibility enables ZIFA-LSNM to achieve a robust representation of the underlying microbial community. We summarize our findings with a discussion and conclusion in Sections 4 and 5, respectively.

## 2. Methods

### 2.1 ZIFA-LSNM model

Let **X**^*n×p*^ *∈* (ℕ ∪ {0})^*n×p*^ denote the *n × p* microbiome count matrix, where *x*_*ij*_ represents the observed read count for taxon *j* in sample *i*. The sequencing depth for the *i*^*th*^ sample is defined as 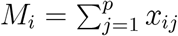, for *i* = 1, …, *n* and *j* = 1, …, *p*. We model the count vector for sample *i*, denoted by ***x***_***i***_ = (*x*_*i*1_, …, *x*_*ip*_)^*T*^ as following a multinomial distribution with total count *M*_*i*_ and probability vector ***ρ***_***i***_ = (*ρ*_*i*1_, …, *ρ*_*ip*_)^*T*^ . Note that this probability vector ***ρ***_***i***_ represents unobserved proportions and is an element of the (*p −* 1) - simplex ***S***^*p−*1^, satisfying 0 *< ρ*_*ij*_ *<* 1 and 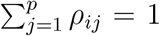. In this article, we map ***ρ***_***i***_ from the simplex to an unconstrained real space using the ALR transformation. Let ***a***_***i***_ = (*a*_*i*1_, …, *a*_*i*(*p−* 1)_)^*T*^ *∈* ℝ^*p−*1^ be the transformed vector for sample *i*. Using the *p*th taxon as the reference, the ALR transformation is defined as:

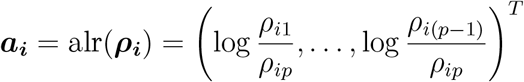

This ALR transformation is bijective. Hence, the original proportions can be recovered through its inverse,

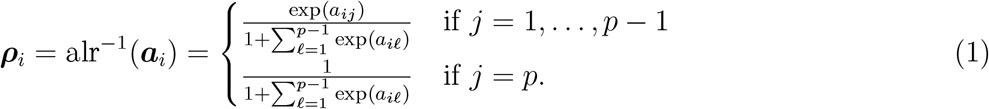

Due to their compositional nature, raw microbiome data cannot be directly analyzed using standard factor analysis, which requires data to reside in a Euclidean space. To appropriately apply a (Bayesian) factor analysis framework, we first map the compositional vector ***ρ***_***i***_ from the simplex to ℝ^*p−*1^ using the ALR transformation. We then model each element of the transformed vector ***a***_***i***_ as a linear combination of a *k*-dimensional vector of latent factors ***F***_*i*_ = (*F*_*i*1_, …, *F*_*ik*_)^*T*^ :

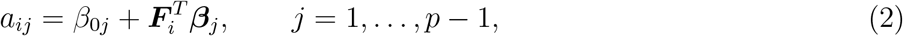

where *β*_0*j*_ represents the taxon-specific intercept and ***β***_***j***_ = (*β*_*j*1_, …, *β*_*jk*_)^*T*^ is the corresponding *k −*dimensional vector of factor loadings. Because the ALR representation is (*p −*1)-dimensional, we adopt the convention *β*_0*p*_ = 0 and ***β***_*p*_ = **0**, so that *a*_*ip*_ ≡0 in (1). We further place priors on the latent factors and loadings. Specifically, we assume a skew-normal distribution for each latent factor *F*_*it*_ and a normal prior for each factor loading *β*_*jt*_.

To account for the excess of zeros in microbiome data, we introduce a latent binary variable, *z*_*ij*_, for *j* = 1, …, *p* − 1. This variable follows a taxon-specific Bernoulli distribution, i.e., *z*_*ij*_*∼* ℬern(*κ*_*j*_) where *κ*_*j*_ is the probability of zero-inflation for taxon *j*. When *z*_*ij*_ = 1, which occurs with probability *κ*_*j*_, the *j*th component of the composition is set to zero. Also, we implicitly assume that *x*_*ij*_ = 0 if *ρ*_*ij*_ = 0.

For *i* = 1, …, *n, j* = 1, …, *p−* 1, *t* = 1, …, *k*, we propose the Bayesian ZIFA-LSNM model in which the likelihood is given by:

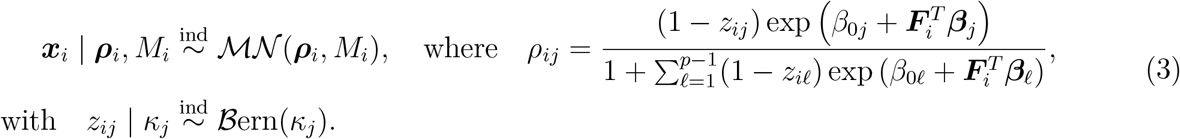

Here, ℳ*N*and ℬern denote multinomial and Bernoulli distributions. Note that the ALR transformation requires *ρ*_*ip*_ *>* 0 and this is a property of the ALR transformation rather than a restriction imposed on the data. Moreover, this restricts only the underlying composition. The reference taxon may still have observed zero counts *x*_*ip*_ = 0. Also, the sequencing depth 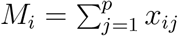 continues to be computed over all *p* taxa. We complete the model definition with priors on the latent factors and the remaining parameters,

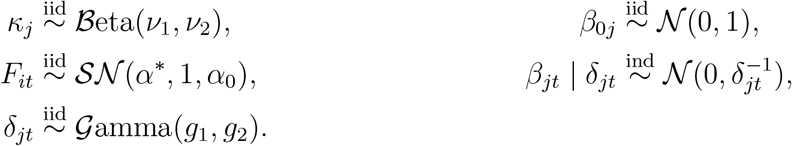

where *N* and *SN*denote normal and skew-normal distributions. The vectors ***F***_*i*_ represent unobserved environmental factors or latent scores and ***β***_***j***_ denote the corresponding factor loadings, where *k* is the number of latent factors. The notation *SN*(*α*^*∗*^, 1, *α*_0_) refers to the skew normal distribution with location, scale, and shape parameters given by *α*^*∗*^, 1, and *α*_0_ respectively, where

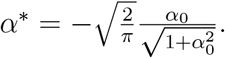

The location parameter *α*^*∗*^ is chosen such that E [*F*_*it*_] = 0, consistent with the assumption of zero-mean latent factors in classical factor analysis. The key innovation is that we use skew-normal priors on the latent factors to explicitly model ALR skewness. In addition, we use informative normal-gamma shrinkage priors on each element of the factor loadings *β*_*jt*_ where *δ*_*jt*_ is the local shrinkage parameter.

The hierarchical architecture of the ZIFA-LSNM model, characterized by multiple layers of latent variables and non-conjugate components, results in an analytically intractable posterior distribution. While MCMC is a gold standard for asymptotically exact inference (Robert and Casella [2004]), its computational burden is often prohibitive for the high-dimensional datasets typical in microbiome research. Thus, to ensure our method is both scalable and efficient, we adopt variational inference, which reframes the challenge of posterior approximation as an optimization problem.

### 2.2 Variational Inference

Let ***Θ*** = (***B, Δ, κ, β***_0_, ***F***, ***z***) be the set of all model parameters and latent variables where, ***B*** = {*β*_*jt*_}_1*≤ j ≤* (*p −* 1), 1 *≤ t ≤ k*_, ***Δ***= {*δ*_*jt*_}_1*≤ j ≤* (*p −* 1), 1 *≤ t ≤ k*_, ***κ*** = {*κ*_*j*_}_1≤ *j* ≤ *p* − 1_, ***β***_0_ = {*β*_0*j*_} _1*≤ j ≤* (*p −* 1)_, ***F*** = {*F*_*it*_} _1 *≤ i≤ n*, 1*≤ t≤ k*_, and ***z*** = {*z*_*ij*_}_1 *≤ i ≤ n*, 1*≤ j ≤* (*p −*1)_. The posterior *p*(***Θ*** |***X***) is analytically intractable because the marginal likelihood,

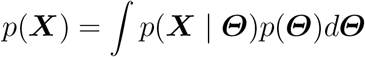

has no closed-form solution.

Variational inference provides a computational alternative to MCMC. It reframes the inference problem as an optimization problem, where the posterior distribution *p*(***Θ*** | ***X***) is approximated by a tractable variational distribution *q*(***Θ***) belonging to some known family of densities *F* (Bishop and Nasrabadi [2006], Ormerod and Wand [2010], Blei et al. [2017]). The best approximation, *q*^*∗*^ *∈ F*, is found by minimizing the reverse Kullback-Leibler (rKL) divergence between *q*(***Θ***) and *p*(***Θ*** | ***X***),

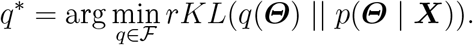

where,

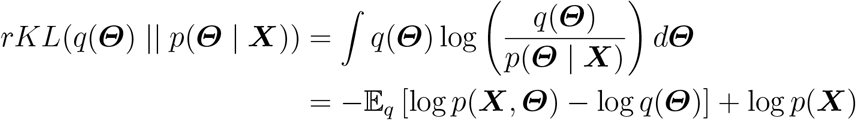

It is clear that minimizing the rKL-divergence is equivalent to maximizing the evidence lower bound (ELBO) where

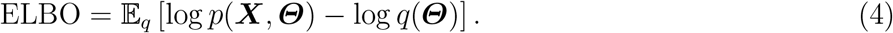

The complexity of the variational family dictates the difficulty of the optimization. To ensure a tractable inference procedure, we adopt the mean-field approximation, which assumes the variational distribution *q*(***Θ***) fully factorizes over all parameters and latent variables as follows:

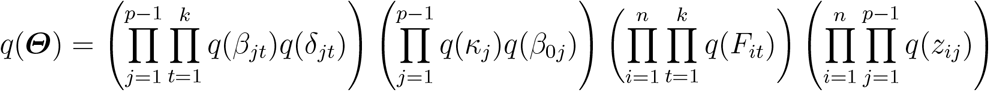

We specify a parametric form for each parameter in the variational family by selecting distributions that balance natural conjugacy with computational convenience. Specifically, we define,

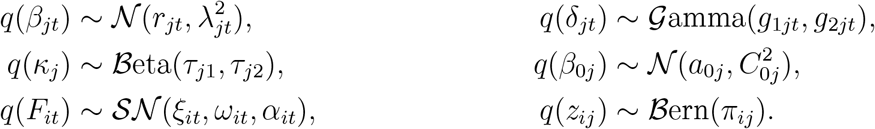

The goal of the variational inference procedure is then to optimize the set of variational parameters,

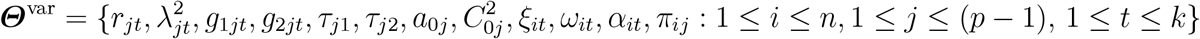

Calculating the ELBO in (4) requires computing the expectations E_*q*_ [log *p*(***X, Θ***)] and E_*q*_ [log *q*(***Θ***)], which becomes analytically challenging. Therefore, we use second order Taylor approximations to evaluate these expressions,

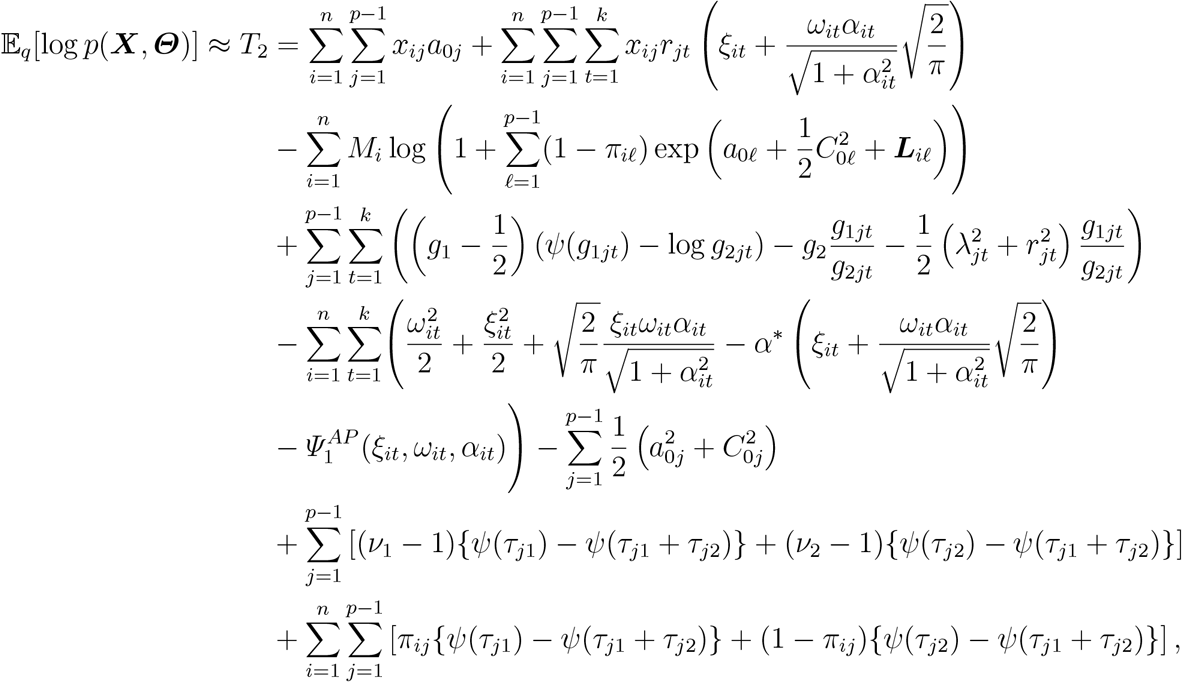

and

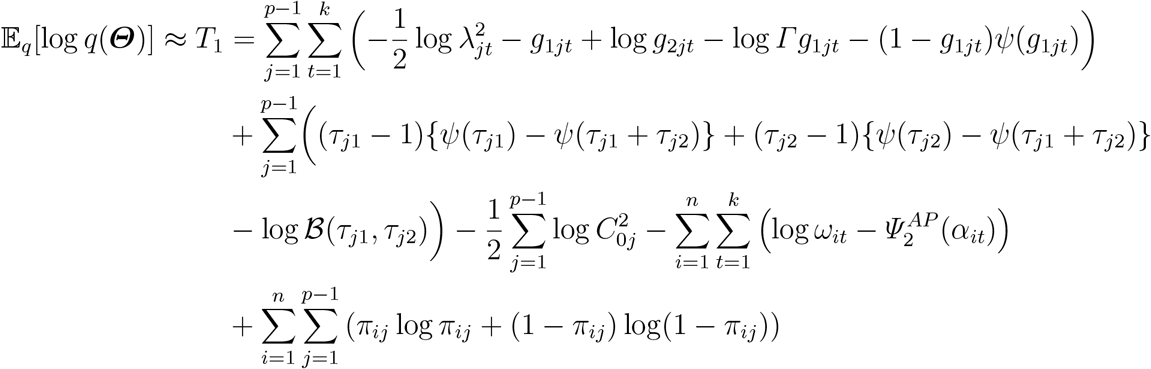

where, ℬ (, ) is the beta function, *ψ* is the digamma function, and

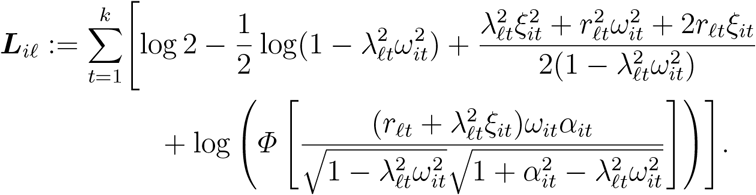

Thus,

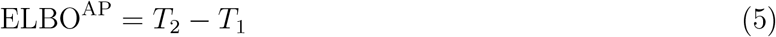

which is a surrogate of the ELBO.

Here and throughout the paper, we drop all constants that do not depend on the parameters. The term ***L***_*il*_ is the sum over *t* = 1 to *k* of the logarithm of the moment generating function of the product of a normal and a skew-normal random variable. From Supplementary Material Section 2, Lemma 2 yields ***L***_*il*_ under *s* = 1. Further details on the derivations and explanations of ELBO^AP^, ***L***_*il*_, 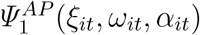, and 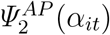 can be found in Supplementary Material. Maximizing ELBO^AP^ with respect to ***Θ***^var^ therefore forms the basis of our inference algorithm.

### 2.3 Parameter estimation

Direct maximization of (5) is difficult due to parameter interactions and the sum of the terms inside the logarithm. We use a variational algorithm that maximizes (5) by alternating updates of {*π*_*ij*_} and ***Θ***^var^ \ {*π*_*ij*_} in turn, and iterating these steps until convergence of ELBO^AP^. This is outlined in Algorithm 1 below. We use singular value decomposition (SVD) initialization for *r*_*jt*_ and *ξ*_*it*_.

#### Algorithm 1

Variational Algorithm for ZIFA-LSNM

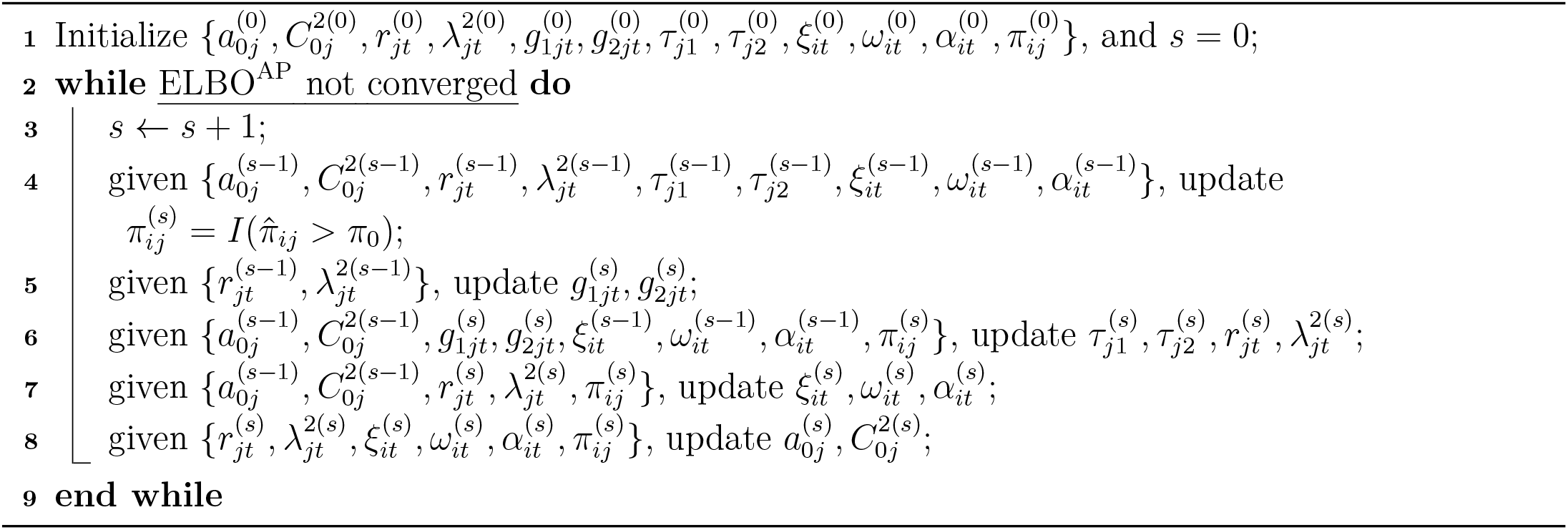

Note that, in total, the number of variational parameters is 4(*p−* 1)*k* + 4(*p −*1) + 3*nk* + *n*(*p −*1), which for *n* = 100, *p* = 50 and *k* = 5 is 7576. Detailed derivations for updating the variational parameters are provided in the Supplementary Material. We obtain the final point estimates for the model parameters by taking the posterior mean of their respective variational distributions,

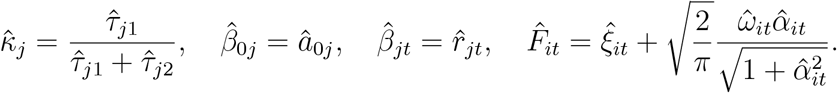

In principle, one could use any statistic from the posterior. We use posterior mean for all point estimation in this paper. Now, given these estimates, the underlying compositions *ρ*_*ij*_ are then estimated as,

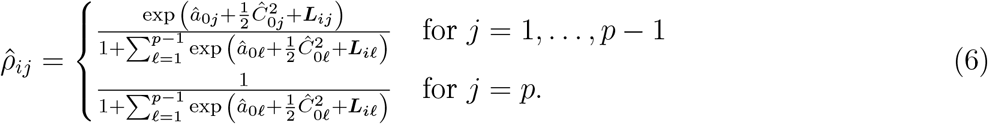

We declare convergence when the relative change of ELBO^AP^ is

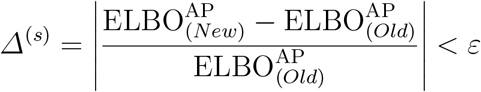

for some small *ε >* 0. Now, given the analytical form of ELBO^AP^ in equation (5), it becomes notoriously difficult to find the update for *π*_*ij*_, primarily due to the log-of-sum terms. Thus, we employ two computational strategies.

#### 2.3.1 Multinomial-Poisson equivalence

First, we leverage the well-established relationship between the multinomial and Poisson regression models to simplify the update of *π*_*ij*_. As noted by Baker [1994], for every multinomial regression model, one can define an equivalent Poisson regression model by introducing a latent parameter. Thus, we provide an equivalent Zero-Inflated Factor Analysis Logistic Skew-Normal Poisson (ZIFA-LSNP) model with the sample-specific latent parameter, *ϒ*_*i*0_, which has the following update

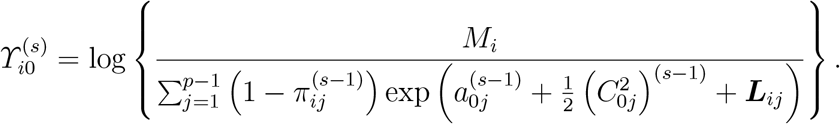

The variational update for *π*_*ij*_ is then consequently derived from this more convenient Poisson-based ELBO,

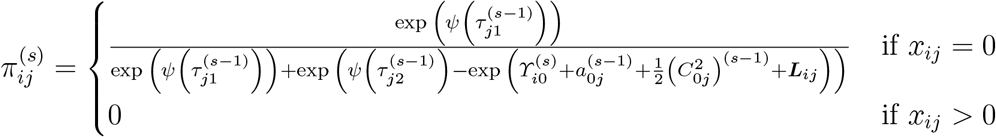

These updates (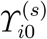 and 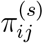) are derived from the ELBO of the ZIFA-LSNP model, which is detailed in the Supplementary Material.

#### 2.3.2 Classification Variational Inference Step

Second, inspired by the classification EM algorithm (Celeux and Govaert [1992]), we introduce a classification step for *π*_*ij*_. In the early stages of optimization, when parameter estimates are unreliable, soft updates of *pi*_*ij*_ can lead to slow or unstable convergence (Celeux and Govaert [1992]). The classification step addresses this by converting it into “hard” assignments (0 or 1) at each iteration,

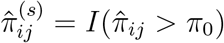

where *π*_0_ is a pre-specified threshold for this classification. Following standard practice in binary classification, we set *π*_0_ = 0.5 as the default value.

### 2.4 Simulation design

The primary objectives of our simulation study are to assess the ZIFA-LSNM model’s ability to: (i) recover the latent structure (loadings and scores); (ii) effectively incorporate zero-inflation and skewness; and (iii) provide reliable estimates of the true underlying microbial compositions.

We compare our model’s performance against ZIPPCA-LPNM [Ba et al., 2024], selected for its improvements over its predecessor ZIPPCA-LNM [Zeng et al., 2023]. Because the factor loadings and latent factors are identified only up to an invertible linear transformation, element-wise error is not a meaningful measure of their recovery. We therefore use the smallest canonical correlation between the true and estimated matrices, i.e., the *k*-th of the *k* canonical correlations. This is the correlation attained by the direction of the true factor space that is least well aligned with the estimated space, and is thus a worst-case summary: it is close to one only if every direction of the space has been recovered, not merely the dominant one. Canonical correlation is widely used as a measure of goodness-of-fit in factor analysis [Bai and Li, 2012, Goyal et al., 2008, Doz et al., 2012]. For the rest of the parameters, element-wise accuracy is the quantity of interest, and we report the root mean squared error (RMSE).

We simulate 250 count datasets for each combination of sample size *n* ∈{50, 100, 1000}, number of taxa *p* ∈{50, 100 }, and true number of factors *k* ∈{3, 5}. For *k* = 3 we consider two levels of latent-factor skewness, *α*_true_ ∈{0, 5} . For *k* = 5, we considered *α*_true_ ∈{0, 3 }. Each simulated dataset is then fitted with the number of factors set to its true value *k* and with three different values of the prior skewness parameter supplied to the algorithm, *α*_0_∈{ 0, 2, 10} . The hyperparameters are *ν*_1_ = 1, *ν*_2_ = 3, *g*_1_ = 1.5, *g*_2_ = 1.5. The specifics of the data-generating process are given in Table 1.

**Table 1:** Specification of the data-generating mechanism for the zero-inflated model.

| Data-generating mechanism for the simulation study |  |  |  |  | Fitted with |
| --- | --- | --- | --- | --- | --- |
| $M_i$ | $\beta_{jt}$ | $\kappa_j$ | $\beta_{0j}$ | $F_{it}$ | $\alpha_0$ |
| $\mathcal{U}(8 \times 10^4, 10^5)$ | $\mathcal{U}(-2.5, 2.5)$ | $\mathcal{U}(0, 0.85)$ | $\mathcal{U}(-2, 2)$ | $\mathcal{SN}(\alpha^*, 1, \alpha_{\text{true}})$ | 0, 2, 10 |

## 3. Results

### 3.1 Skewness simulation

The skewness of an observed ALR coordinate can have two distinct origins. The first is the latent structure: under (2) the ALR transform of the underlying composition is a linear combination of the latent factors, so skewed factors produce skewed *a*_*ij*_. The second is what we call the count layer: zeros generated by the zero-inflation component, multinomial variation, variation in sequencing depth across samples, and a small pseudo-count added before the transformation. Under *α* = 0 there is no latent skewness, so any skewness in the observed ALR is produced by the count layer. Any systematic shift of the observed skewness at *α≠* 0 reflects the asymmetry of the latent factors.

We generate data from the full ZIFA-LSNM model with *n* = 100, *p* = 200 and *k* = 5, setting all loadings to 1 and all intercepts to 0. The zero-inflation probabilities are drawn once from the distibution in Table 1 and held fixed across all values of *α* and all replicates, and the sequencing depths from *U* (8 *×* 10^4^, 10^5^). For each *α ∈ {−*10, *−*5, 0, 5, 10} the latent factors are drawn from *SN* (*α*^*∗*^, 1, *α*) and likewise held fixed across replicates, so that the only quantity differing between the five settings is *α*. In each replicate we add a small pseudo-count *c* to all counts, apply the ALR transformation, and record the sample skewness of each of the *p −*1 coordinates. The experiment is repeated for *c* = 1, 0.5, 10^*−*2^, 10^*−*4^. Results for *c* = 1, 0.5, 10^*−*4^ are given in the Supplementary Material. We note that the addition of pseudo-counts in this way appears to be standard practice (Lin and Peddada [2020], Xia et al. [2013]).

Figure 1 shows, for *c* = 10^*−*2^, that skew-normal factors (*α* ≠ 0) produce more per-taxon skewness in the ALR-transformed counts than Gaussian factors (*α* = 0) under an otherwise identical count and pseudo-count scheme. Thus, in the presence of highly skewed counts, Gaussian latent factors alone may not be sufficient to model this behavior.

**Figure 1:**
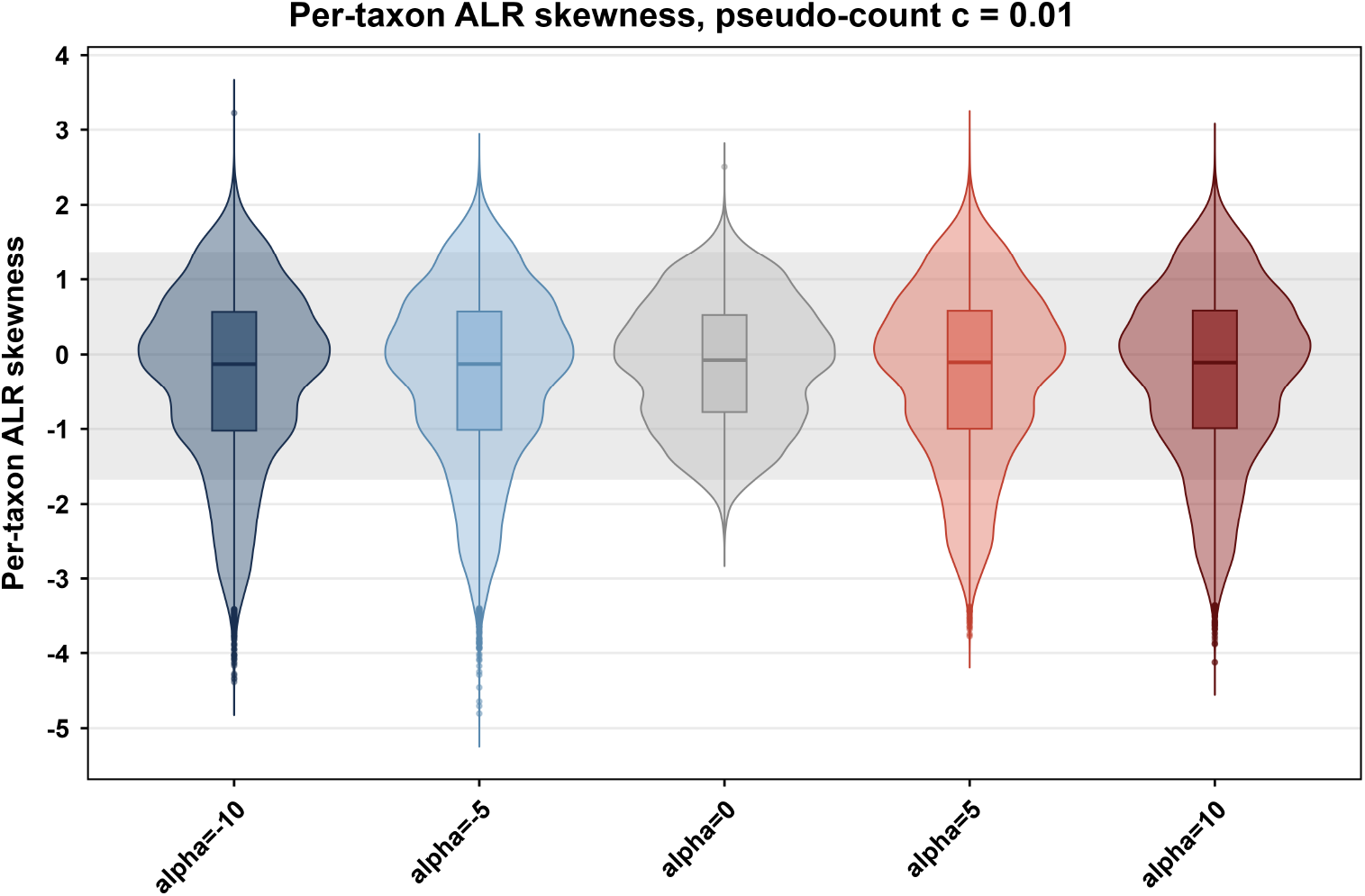
Per-taxon skewness of the ALR-transformed counts after adding a pseudo-count *c* = 10^*−*2^. The gray band is the 95% range of values under *α* = 0.

### 3.2 Model simulation results

Tables 2 and 3 show that ZIFA-LSNM improves on its Gaussian-based counterpart ZIPPCA-LPNM. For the intercepts we report the RMSE of the linear predictor 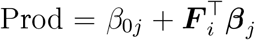rather than of *β*_0*j*_ alone, because

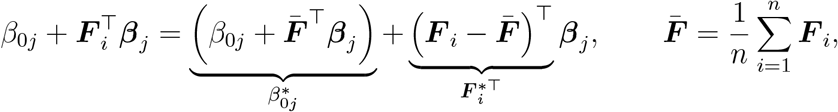

so any shift in the location of the latent factors can be absorbed into the intercept without changing the model. The intercept and the factor mean are therefore not separately identified [Bai and Li, 2012], and the estimation error of *β*_0*j*_ on its own is not a meaningful quantity. Across sample sizes and numbers of taxa, ZIFA-LSNM attains higher canonical correlations for both the latent factors and the factor loadings, and lower RMSEs for *κ*, Prod and ***ρ***. The exceptions are a few settings in which the two models are comparable for the compositions. This advantage holds whether or not the generating distribution is skewed, and the prior skewness *α*_0_ has only a modest influence, which diminishes as the sample size grows.

**Table 2:** Results for *k* = 3. Recovery of the factor loadings ***B*** and the latent factors ***F*** is measured by the smallest non-zero canonical correlation between the true and estimated matrices (larger is better; maximum 1). The remaining quantities are assessed by RMSE (smaller is better; minimum 0).

| $\alpha_{\text{true}}$ | $(n, p)$ | Model | $\alpha_0$ | Canonical Correlation | | RMSE | | |
| --- | --- | --- | --- | --- | --- | --- | --- | --- |
| | | | | $\mathbf{B}$ | $\mathbf{F}$ | $\kappa$ | Prod | $\rho$ |
| 0 | (50, 50) | ZIPPCA-LPNM | — | 0.0729 | 0.1801 | 0.1548 | 2.5177 | 0.0479 |
|  |  | ZIFA-LSNM | 0 | 0.9690 | 0.9979 | 0.0606 | 0.3786 | 0.0101 |
|  |  | ZIFA-LSNM | 2 | 0.9709 | 0.9979 | 0.0606 | 0.3567 | 0.0096 |
|  |  | ZIFA-LSNM | 10 | 0.9829 | 0.9992 | 0.0602 | 0.2130 | 0.0058 |
| 0 | (100, 50) | ZIPPCA-LPNM | — | 0.1355 | 0.1417 | 0.1529 | 2.7793 | 0.0530 |
|  |  | ZIFA-LSNM | 0 | 0.9990 | 0.9994 | 0.0477 | 0.1275 | 0.0014 |
|  |  | ZIFA-LSNM | 2 | 0.9991 | 0.9993 | 0.0478 | 0.1250 | 0.0014 |
|  |  | ZIFA-LSNM | 10 | 0.9995 | 0.9989 | 0.0477 | 0.1148 | 0.0012 |
| 0 | (1000, 50) | ZIPPCA-LPNM | — | 0.0568 | 0.2518 | 0.1377 | 2.7025 | 0.0520 |
|  |  | ZIFA-LSNM | 0 | 0.9999 | 0.9997 | 0.0194 | 0.0546 | 0.0004 |
|  |  | ZIFA-LSNM | 2 | 0.9999 | 0.9998 | 0.0196 | 0.0316 | 0.0004 |
|  |  | ZIFA-LSNM | 10 | 0.9999 | 0.9991 | 0.0196 | 0.0711 | 0.0008 |
| 0 | (50, 100) | ZIPPCA-LPNM | — | 0.0494 | 0.1898 | 0.1310 | 2.7658 | 0.0325 |
|  |  | ZIFA-LSNM | 0 | 0.8865 | 0.9935 | 0.0627 | 0.9079 | 0.0208 |
|  |  | ZIFA-LSNM | 2 | 0.8800 | 0.9954 | 0.0626 | 0.8689 | 0.0211 |
|  |  | ZIFA-LSNM | 10 | 0.9311 | 0.9942 | 0.0626 | 0.5889 | 0.0124 |
| 0 | (100, 100) | ZIPPCA-LPNM | — | 0.0658 | 0.1362 | 0.1196 | 2.5368 | 0.0340 |
|  |  | ZIFA-LSNM | 0 | 0.9881 | 0.9982 | 0.0486 | 0.1994 | 0.0017 |
|  |  | ZIFA-LSNM | 2 | 0.9920 | 0.9986 | 0.0483 | 0.1784 | 0.0014 |
|  |  | ZIFA-LSNM | 10 | 0.9885 | 0.9979 | 0.0483 | 0.2044 | 0.0016 |
| 0 | (1000, 100) | ZIPPCA-LPNM | — | 0.0493 | 0.2126 | 0.1183 | 2.7370 | 0.0307 |
|  |  | ZIFA-LSNM | 0 | 0.9999 | 0.9999 | 0.0295 | 0.0958 | 0.0001 |
|  |  | ZIFA-LSNM | 2 | 0.9999 | 0.9999 | 0.0296 | 0.1048 | 0.0001 |
|  |  | ZIFA-LSNM | 10 | 0.9999 | 0.9996 | 0.0295 | 0.0750 | 0.0002 |
| 5 | (50, 50) | ZIPPCA-LPNM | — | 0.0832 | 0.2683 | 0.0709 | 1.9193 | 0.0352 |
|  |  | ZIFA-LSNM | 0 | 0.9087 | 0.9830 | 0.0611 | 0.4948 | 0.0150 |
|  |  | ZIFA-LSNM | 2 | 0.8943 | 0.9938 | 0.0610 | 0.5618 | 0.0162 |
|  |  | ZIFA-LSNM | 10 | 0.9403 | 0.9930 | 0.0606 | 0.2831 | 0.0086 |
| | | | | $\mathbf{B}$ | $\mathbf{F}$ | $\kappa$ | Prod | $\rho$ |
| 5 | (100, 50) | ZIPPCA-LPNM | — | 0.0738 | 0.1682 | 0.0553 | 1.7777 | 0.0361 |
|  |  | ZIFA-LSNM | 0 | 0.9924 | 0.9987 | 0.0439 | 0.0857 | 0.0012 |
|  |  | ZIFA-LSNM | 2 | 0.9960 | 0.9993 | 0.0439 | 0.0601 | 0.0009 |
|  |  | ZIFA-LSNM | 10 | 0.9998 | 0.9995 | 0.0435 | 0.0530 | 0.0005 |
| 5 | (1000, 50) | ZIPPCA-LPNM | — | 0.0915 | 0.4278 | 0.0793 | 1.7920 | 0.0376 |
|  |  | ZIFA-LSNM | 0 | 0.9999 | 0.9999 | 0.0137 | 0.0256 | 0.0002 |
|  |  | ZIFA-LSNM | 2 | 0.9999 | 0.9999 | 0.0137 | 0.0263 | 0.0002 |
|  |  | ZIFA-LSNM | 10 | 0.9999 | 0.9999 | 0.0136 | 0.0276 | 0.0003 |
| 5 | (50, 100) | ZIPPCA-LPNM | — | 0.0523 | 0.3004 | 0.0685 | 2.0144 | 0.0198 |
|  |  | ZIFA-LSNM | 0 | 0.9822 | 0.9978 | 0.0603 | 0.1548 | 0.0028 |
|  |  | ZIFA-LSNM | 2 | 0.9822 | 0.9978 | 0.0604 | 0.1465 | 0.0022 |
|  |  | ZIFA-LSNM | 10 | 0.9850 | 0.9977 | 0.0599 | 0.1128 | 0.0019 |
| 5 | (100, 100) | ZIPPCA-LPNM | — | 0.0743 | 0.1821 | 0.0661 | 1.7836 | 0.0216 |
|  |  | ZIFA-LSNM | 0 | 0.9994 | 0.9998 | 0.0432 | 0.0648 | 0.0005 |
|  |  | ZIFA-LSNM | 2 | 0.9994 | 0.9998 | 0.0428 | 0.0798 | 0.0004 |
|  |  | ZIFA-LSNM | 10 | 0.9997 | 0.9994 | 0.0432 | 0.0705 | 0.0004 |
| 5 | (1000, 100) | ZIPPCA-LPNM | — | 0.0430 | 0.1866 | 0.0503 | 1.9476 | 0.0206 |
|  |  | ZIFA-LSNM | 0 | 0.9999 | 0.9999 | 0.0140 | 0.0364 | 0.0001 |
|  |  | ZIFA-LSNM | 2 | 0.9999 | 0.9999 | 0.0139 | 0.0404 | 0.0001 |
|  |  | ZIFA-LSNM | 10 | 0.9999 | 0.9999 | 0.0139 | 0.0351 | 0.0001 |

**Table 3:**
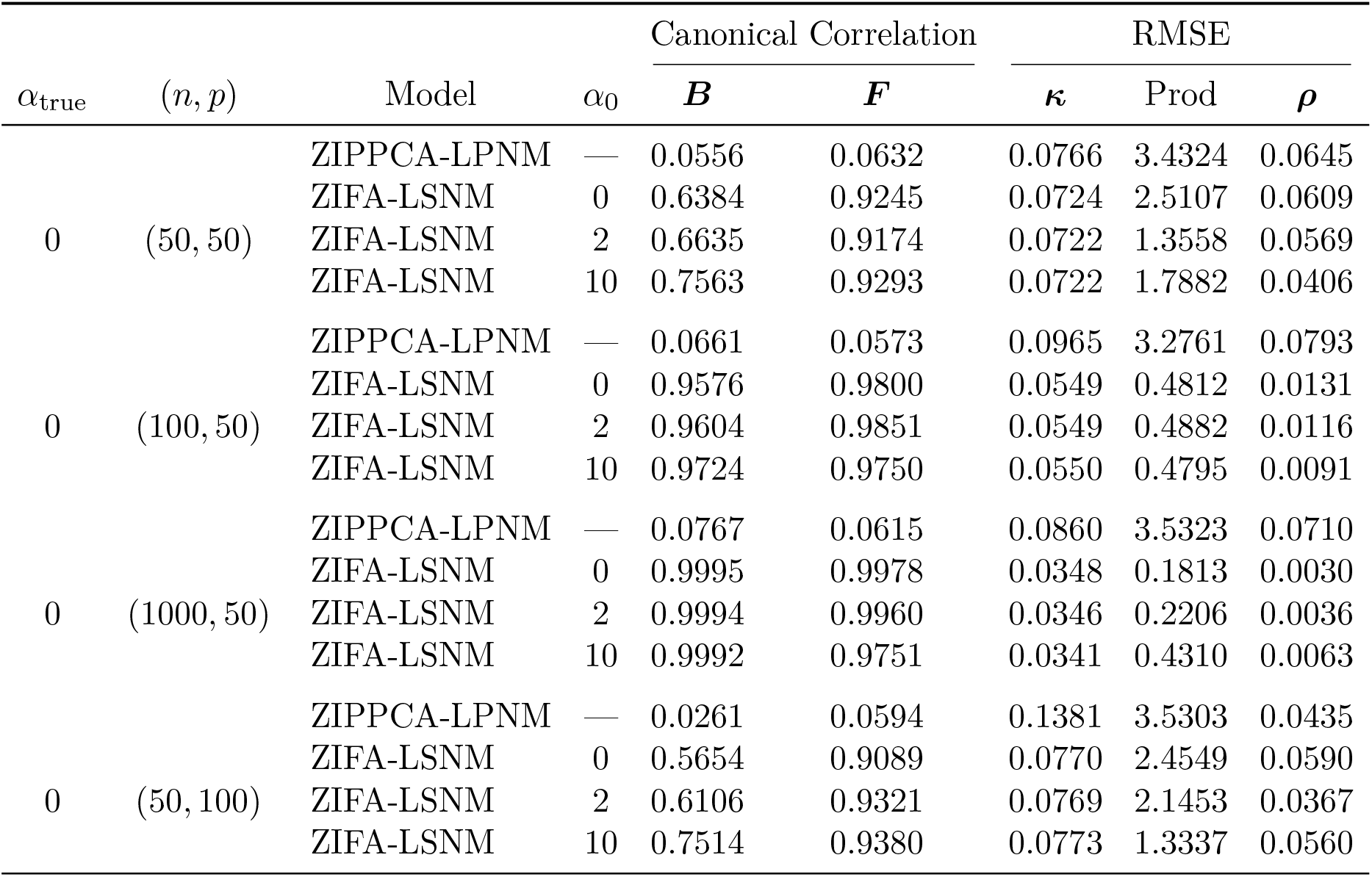
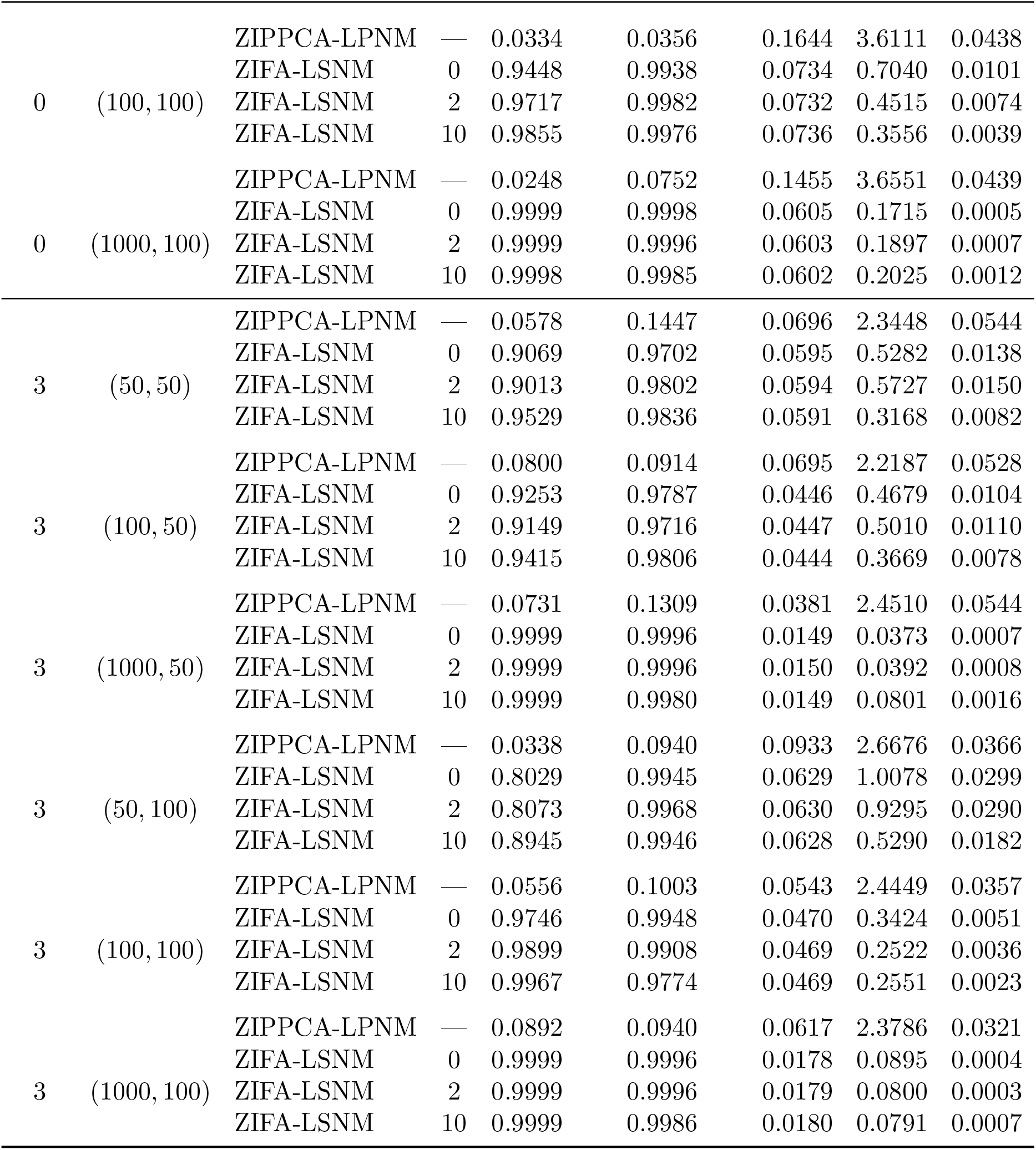
Results for *k* = 5. Recovery of the factor loadings ***B*** and the latent factors ***F*** is measured by the smallest non-zero canonical correlation between the true and estimated matrices (larger is better; maximum 1). The remaining quantities are assessed by RMSE (smaller is better; minimum 0).

ZIFA-LSNM recovers the latent structure (latent factors and factor loadings) and the compositions in most configurations, and its accuracy improves with sample size in almost every case. Across all 24 combinations, the RMSEs of ***κ*** and ***ρ*** decrease with increase in *n*. The RMSE of Prod does so in 23 of 24 and the canonical correlation for ***B*** in 23 of 24. The exception in both cases is the configuration *k* = 5, *α*_true_ = 3, *p* = 50. Comparing *n* = 50 with *n* = 1000, all four (***B, κ***, Prod, ***ρ***) quantities improve in all 24 configurations. The latent factors are the most robustly recovered quantity, attaining a canonical correlation of at least 0.90 in every configuration, while the loadings and the compositions are the most sensitive to small samples. By contrast, ZIPPCA-LPNM does not recover the latent structure in any configuration: its canonical correlations never exceed 0.14 for ***B*** or 0.43 for ***F*** .

The two tables differ in how much the sample size matters. With *k* = 3 (Table 2) ZIFA-LSNM performs well at every sample size, and is more accurate than ZIPPCA-LPNM on every measure in every row. With *k* = 5 (Table 3) this continues to hold for *n* = 100 and *n* = 1000, but at *n* = 50 the outcome depends on *α*_true_. With skewed factors (*α*_true_ = 3) recovery is reduced but still acceptable. With Gaussian factors (*α*_true_ = 0) the weakest loading direction is poorly recovered for both *p* = 50 and *p* = 100, and at *p* = 100 the compositions suffer as well. This is the only part of the table in which the advantage of ZIFA-LSNM is small or, for the compositions, absent. Even there the factor scores are recovered far more accurately than by ZIPPCA-LPNM, and raising the sample size to *n* = 100 removes the deficit entirely.

In summary, ZIFA-LSNM recovered the latent factors reliably in every configuration, and recovered the loadings and compositions accurately in all but the *n* = 50 settings with *k* = 5. The advantage over ZIPPCA-LPNM was present whether the generating factors were skewed (*α*_true_ ≠ 0) or Gaussian (*α*_true_ = 0), indicating that the skew-normal prior does not degrade performance when the factors are Gaussian. Across the range of asymmetry considered, ZIFA-LSNM therefore provides a more faithful representation of the microbial community than ZIPPCA-LPNM.

### 3.3 Data application and results

In this section, we apply the ZIFA-LSNM model to a publicly available dataset from a family-based study of inflammatory bowel disease (IBD) (Muller et al. [2022], Jacobs et al. [2016]). The dataset comprises 16S rRNA sequencing data from 90 participants: 54 healthy controls and 36 IBD patients (26 with Crohn’s disease [CD] and 10 with ulcerative colitis [UC]).

The raw count data were aggregated to the genus level, with unclassified taxa removed. To reduce extreme sparsity, we applied two standard prevalence filters: genera were retained if the proportion of zeros was below 85%, and if the cumulative read count exceeded 20. The filtered dataset comprised 132 genera across 90 samples.

Table 4 and Figures 2(a) and 2(b) report the in-sample area under the curve (AUC) and the crossvalidated AUC (CV-AUC) obtained when the varimax-rotated posterior mean factor scores are used as predictors in a logistic regression for the binary outcome (IBD versus healthy controls). CV-AUC was computed by 5-fold cross-validation, and the reported value is the mean over 100 independent random partitions. ZIFA-LSNM outperforms ZIPPCA-LPNM at every value of *k* under both metrics: the CV-AUC of ZIPPCA-LPNM remains between 0.461 and 0.536 for *k≤* 5, and although it improves for *k ≥* 6 it never reaches the ZIFA-LSNM value at the same *k*. For ZIFA-LSNM the CV-AUC rises up to *k* = 5, gains little thereafter, and declines for most *α*_0_ from *k* = 8 onwards, indicating that further factors capture noise rather than structure. The gain from *k* = 5 to *k* = 7 is smaller than the spread across the four values of *α*_0_ at *k* = 7 and the difference in CV-AUC between *k* = 5 and *k* = 6 is small relative to the variation across *α*_0_, so parsimony governs the choice. We therefore select *k* = 5, the smallest value in this flat region whose CV-AUC also changes little across the choice of prior.

**Table 4:** In-sample and cross-validated AUC for ZIPPCA-LPNM and ZIFA-LSNM across the number of latent factors *k*. Cross-validated AUC is the mean over 100 random 5-fold partitions.

| $k$ | Model | $\alpha_0$ | AUC | CV-AUC | $k$ | Model | $\alpha_0$ | AUC | CV-AUC |
| --- | --- | --- | --- | --- | --- | --- | --- | --- | --- |
| 2 | ZIPPCA-LPNM | — | 0.6167 | 0.5363 | 7 | ZIPPCA-LPNM | — | 0.8292 | 0.7176 |
|  | ZIFA-LSNM | 2 | 0.7782 | 0.7335 |  | ZIFA-LSNM | 2 | 0.8492 | 0.7601 |
|  | ZIFA-LSNM | −2 | 0.7299 | 0.6719 |  | ZIFA-LSNM | −2 | 0.8364 | 0.7435 |
|  | ZIFA-LSNM | 5 | 0.7731 | 0.7343 |  | ZIFA-LSNM | 5 | 0.8806 | 0.7982 |
|  | ZIFA-LSNM | −5 | 0.7289 | 0.6714 |  | ZIFA-LSNM | −5 | 0.8425 | 0.7516 |
| 3 | ZIPPCA-LPNM | — | 0.5910 | 0.4606 | 8 | ZIPPCA-LPNM | — | 0.8132 | 0.7015 |
|  | ZIFA-LSNM | 2 | 0.7752 | 0.7306 |  | ZIFA-LSNM | 2 | 0.8405 | 0.7464 |
|  | ZIFA-LSNM | −2 | 0.7839 | 0.7367 |  | ZIFA-LSNM | −2 | 0.8590 | 0.7524 |
|  | ZIFA-LSNM | 5 | 0.7782 | 0.7307 |  | ZIFA-LSNM | 5 | 0.8317 | 0.7377 |
|  | ZIFA-LSNM | −5 | 0.7736 | 0.7278 |  | ZIFA-LSNM | −5 | 0.8585 | 0.7707 |
| 4 | ZIPPCA-LPNM | — | 0.6476 | 0.5214 | 9 | ZIPPCA-LPNM | — | 0.8055 | 0.6734 |
|  | ZIFA-LSNM | 2 | 0.7926 | 0.7377 |  | ZIFA-LSNM | 2 | 0.8266 | 0.7071 |
|  | ZIFA-LSNM | −2 | 0.7716 | 0.7075 |  | ZIFA-LSNM | −2 | 0.8533 | 0.7340 |
|  | ZIFA-LSNM | 5 | 0.8009 | 0.7509 |  | ZIFA-LSNM | 5 | 0.8230 | 0.6968 |
|  | ZIFA-LSNM | −5 | 0.7890 | 0.7303 |  | ZIFA-LSNM | −5 | 0.8580 | 0.7316 |
| 5 | ZIPPCA-LPNM | — | 0.6270 | 0.4726 | 10 | ZIPPCA-LPNM | — | 0.7793 | 0.6168 |
|  | ZIFA-LSNM | 2 | 0.8158 | 0.7466 |  | ZIFA-LSNM | 2 | 0.8564 | 0.7227 |
|  | ZIFA-LSNM | −2 | 0.8220 | 0.7534 |  | ZIFA-LSNM | −2 | 0.8605 | 0.7168 |
|  | ZIFA-LSNM | 5 | 0.8215 | 0.7530 |  | ZIFA-LSNM | 5 | 0.8251 | 0.6963 |
|  | ZIFA-LSNM | −5 | 0.8194 | 0.7469 |  | ZIFA-LSNM | −5 | 0.8580 | 0.7304 |
| 6 | ZIPPCA-LPNM | — | 0.7762 | 0.6783 |  |  |  |  |  |
|  | ZIFA-LSNM | 2 | 0.8281 | 0.7492 |  |  |  |  |  |
|  | ZIFA-LSNM | −2 | 0.8323 | 0.7606 |  |  |  |  |  |
|  | ZIFA-LSNM | 5 | 0.8256 | 0.7558 |  |  |  |  |  |
|  | ZIFA-LSNM | −5 | 0.8307 | 0.7603 |  |  |  |  |  |

**Figure 2:**
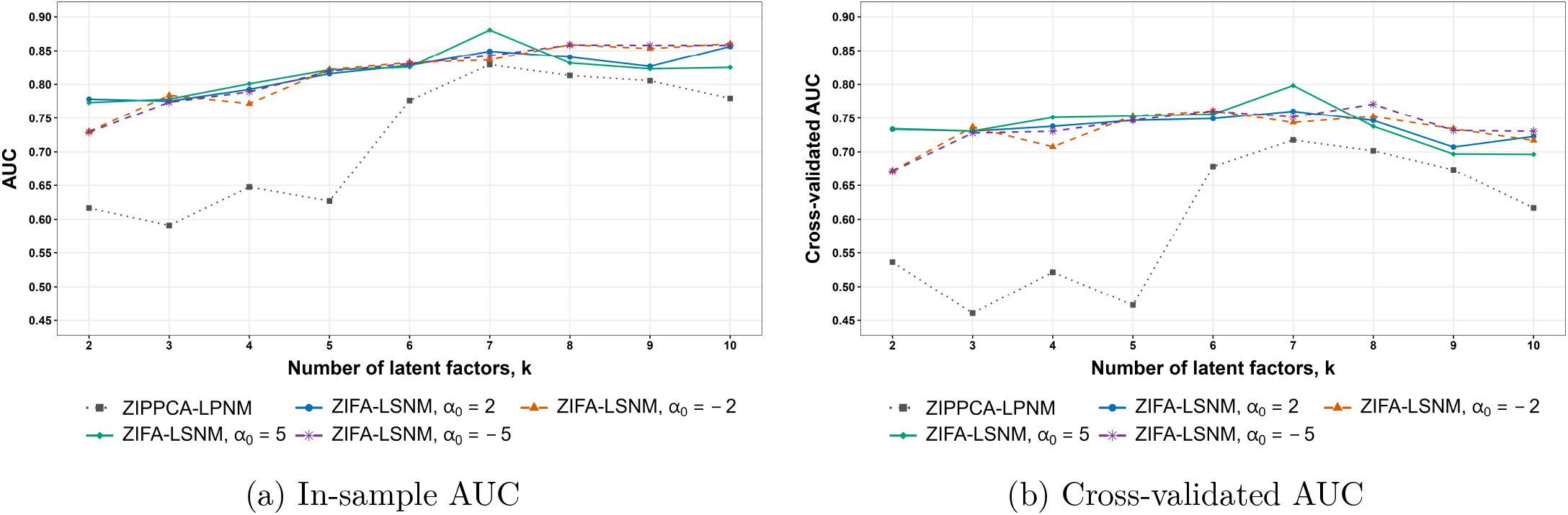
Discrimination of IBD from healthy controls by logistic regression on the estimated factor scores, plotted against the number of latent factors *k*, for ZIPPCA-LPNM and ZIFA-LSNM (*α*_0_ *∈ {±*2, *±*5*}*)

Had the number of latent factors instead been chosen from the ZIPPCA-LPNM fits, the CV-AUC would have been maximized at *k* = 7. This is consistent with the observation of Tu et al. [2024] and Lin et al. [2007] that the Gaussian distribution tends to overestimate the number of components by using additional components to capture the skewness.

The pairwise scatter plots of estimated factor scores for *k* = 5 are shown in Figure 3. To quantify the separation between diseased and healthy individuals in this five-dimensional latent space, we report the Mahalanobis distance (*D*^2^) between group centroids, computed with the pooled within-group covariance, together with an *F* -test derived from the two-sample Hotelling’s *T* ^2^ statistic. This approach has been used to quantify cluster separation in principal component analysis [Goodpaster and Kennedy, 2011]. We apply it here to the factor scores obtained from our proposed model. Results are given in Table 5, which shows a clear ordering across the three comparisons. Crohn’s disease is the most strongly separated from healthy individuals. The pooled IBD group remains highly significant but is attenuated, which is the expected consequence of combining Crohn’s disease with ulcerative colitis samples that are themselves not separable from healthy individuals.

**Table 5:** Separation between disease groups and healthy individuals for *k* = 5 factor scores. *D*^2^ is the Mahalanobis distance between group centroids computed with the pooled within-group covariance. The *F* -statistic is obtained from the two-sample Hotelling’s *T* ^2^.

| Comparison | $n_{\text{Healthy}}$ | $n_{\text{case}}$ | $D^2$ | $F$ | p-value |
| --- | --- | --- | --- | --- | --- |
| CD vs Healthy | 54 | 26 | 2.8859 | 9.6099 | $4.25 \times 10^{-7}$ |
| (CD+UC) vs Healthy | 54 | 36 | 1.6776 | 6.9178 | $1.88 \times 10^{-5}$ |
| UC vs Healthy | 54 | 10 | 1.1093 | 1.7511 | 0.1374 |

**Figure 3:**
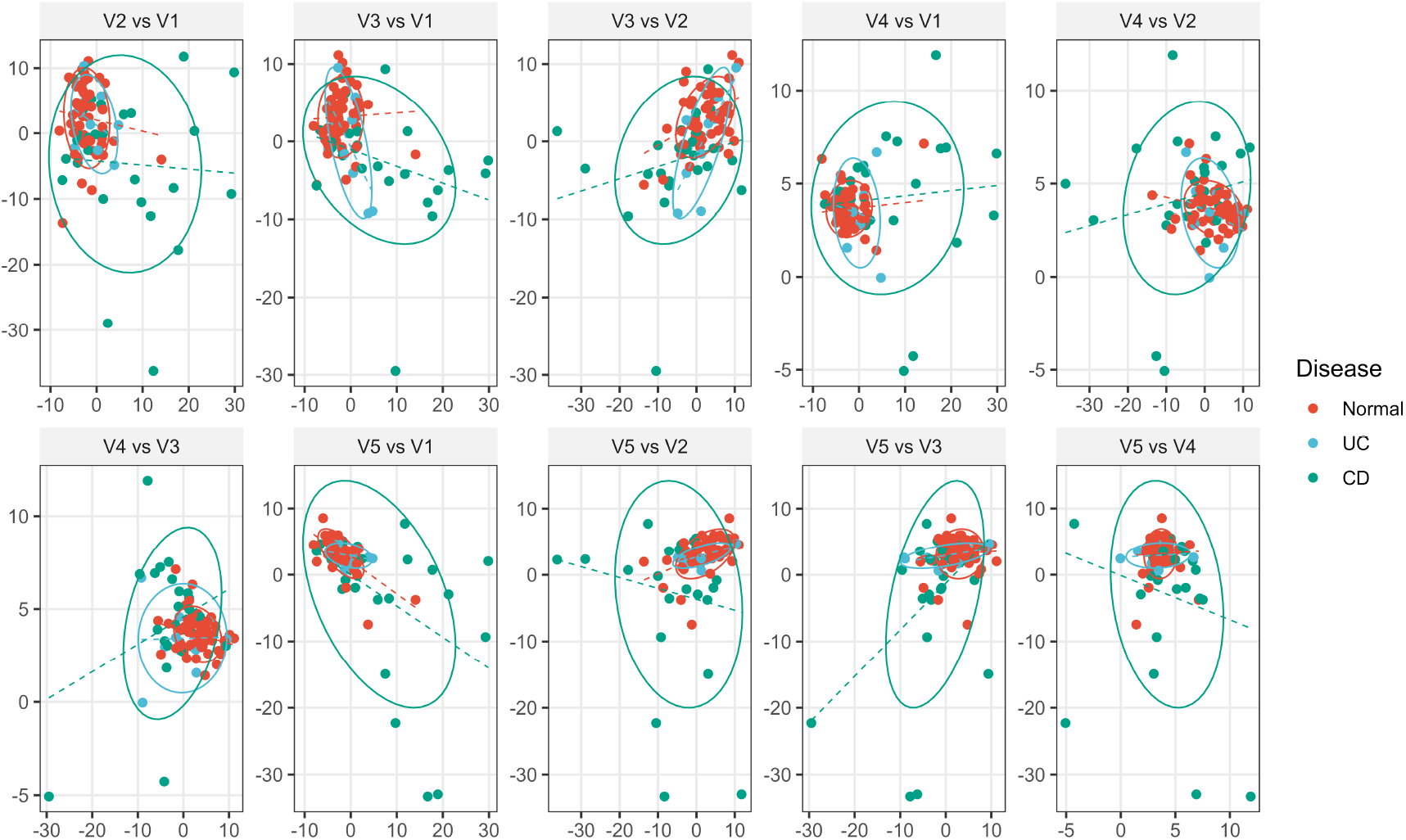
Latent Microbial Factors Distinguish Healthy Controls from IBD Patients. Pairwise scatter plots of factor scores from the 5 latent variables via ZIFA-LSNM model. Each point corresponds to an individual sample and is colored by diagnosis. V*t* denotes factor *t*.

## 4. Discussion

The analysis of microbiome data presents a unique set of statistical challenges, primarily the need to achieve effective dimension reduction within a framework that rigorously accounts for both compositionality and zero inflation. While progress has been made, the prevalent issue of skewness in log-ratio-transformed compositions has been largely overlooked. This paper introduces the ZIFA-LSNM model, a Bayesian hierarchical framework designed to handle these challenges while accounting for skewness through the use of skew-normal priors on the latent factors.

Our simulation studies provide evidence of improved performance of ZIFA-LSNM compared to its Gaussian-based counterpart. Across a range of scenarios, our model demonstrated higher canonical correlations for the latent structure and lower RMSEs for ***κ***, Prod and, ***ρ*** in all but a few small-sample settings. These findings suggest that explicitly modeling skewness can enhance both the statistical robustness and biological interpretability of microbiome data. Furthermore, our adoption of a variational inference framework ensures computational scalability, making the approach feasible for modern, large-scale microbiome studies.

Our approach uses variational inference to find the estimates. The criterion optimized by Algorithm 1, ELBO^AP^, is a surrogate of the ELBO in which two terms are second order Taylor approximations. In all simulated and real datasets we fitted, ELBO^AP^ increased monotonically. As with any variational algorithm, convergence is to a local rather than a global optimum. There are multiple strategies that could be explored to avoid getting stuck in local optima. First, one may utilize multiple random initializations for different variational parameters (Bishop and Nasrabadi [2006], Blei et al. [2017]). Alternatively, initializing variational parameters for factor loadings and factor scores using a low-rank decomposition such as PCA or singular value decomposition (SVD) often improves convergence and reduces sensitivity to poor local optima (Tipping and Bishop [1999]). Here, we have used the SVD initialization approach to address the potential convergence issue.

While the proposed model is effective, it possesses limitations that highlight opportunities for future research. Although our variational inference framework is scalable, its computational complexity can be substantial when fitting count-based models to high-dimensional microbiome data with large sample sizes (*n*) and many taxa (*p*). This can present challenges in ultra-high-dimensional settings. Our local shrinkage priors provide effective regularization, but more advanced schemes could be explored. For instance, incorporating Gamma process global-local shrinkage priors (Bhattacharya and Dunson [2011], Hansen et al. [2025]), could further enhance factor recovery. Moreover, in the absence of prior knowledge, more sophisticated selection methods can be used to find *k*, such as information criteria like the Variational BIC (You et al. [2014]) that allow *k* to be inferred directly from the data. In addition, a well-known shortcoming of the mean field approximation is that it typically underestimates uncertainty MacKay [2003], Wang and Titterington [2005]). This behavior arises directly from the independence assumption of the mean field approximation as the covariance contributions between the parameters are ignored. By using the sensitivity of the mean field posterior estimates, Giordano et al. [2018] developed a methodology that augments mean field estimates to deliver improved covariance estimates for the model parameters. This technique can be used to provide an improved estimator for the exact marginal covariance of the model parameters.

## 5. Conclusions

Our simulation and real data results indicate that ZIFA-LSNM improves on its Gaussian latent-factor counterpart, ZIPPCA-LPNM, in the presence of large skewness in the ALR-transformed microbiome data. In such data, Gaussian latent factors known to overestimate the number of components because they use additional components to capture skewness (Lin et al. [2007]). Our data application (Section 3.3) is consistent with this: ZIFA-LSNM required fewer factors (*k* = 5) than ZIPPCA-LPNM (*k* = 7).

## 6. Declarations

### Ethics approval and consent to participate

Not applicable.

### Consent for publication

Not applicable.

### Availability of data and materials

The proposed method, `zifalsnm`, is implemented in an openly available R package and it is available at: https://github.com/SaurabhP-MS/zifalsnm/. All reproducible R code, real-data analyses, and simulation results are available at: https://github.com/SaurabhP-MS/zifalsnm-reproducibility/tree/main/Reproducible_R_Codes. The real microbiome dataset analyzed in this study was obtained from the publicly available curated microbiome-metabolome data repository at, https://github.com/borenstein-lab/microbiome-metabolome-curated-data, where in the folder name “data/processed data” it is listed under the dataset name `JACOBS IBD FAMILIES 2016`. The original article associated with this dataset is Jacobs et al. [2016]. The accession number for the dataset is PRJNA324147, and all required articles have been cited in this manuscript.

### Authors’ contributions

SP contributed in methodological theory and application, coded, wrote an R package, conducted simulation studies, created and synthesized all tables and figures, performed real data analysis, primary manuscript preparation; HJ & KM contributed in methodological theory and application, supervised, manuscript preparation. All authors read and approved the final manuscript.

## Acknowledgements

Not applicable.

