## Supplementary Material for "Zero-inflated Bayesian factor analysis model with skew-normal priors for modeling microbiome data"

#### Contents

**1 Derivation of ELBO$^{AP}$** **2**

1.1 Derivation of $\mathbb{E}_q[\log q(\boldsymbol{\Theta})]$ . . . . . 2

1.2 Derivation of $\mathbb{E}_q[\log p(\boldsymbol{x}, \boldsymbol{\Theta})]$ . . . . . 3

**2 MGF of Product of Normal and Skew Normal Random Variables** **6**

**3 Parameter Updates** **8**

**4 Derivatives of $L_{ij}$ Function** **14**

**5 Taylor Expansion and it's gradients for $\Psi_1(\xi_{it}, \omega_{it}, \alpha_{it})$** **15**

**6 Taylor Expansion and it's gradients for $\Psi_2(\alpha_{it})$** **17**

**7 Update for $\pi_{ij}$ parameter** **18**

**8 Skewness Plots** **21**

8.1 Simulation plots . . . . . 21

8.2 Real data plots . . . . . 21

### 1 Derivation of ELBO$^{\mathrm{AP}}$

We split the derivation of ELBO$^{\mathrm{AP}}$ into two parts. Throughout the Supplementary Material, we drop all constants that do not depend on the parameters.

#### 1.1 Derivation of $\mathbb{E}_q[\log q(\boldsymbol{\Theta})]$

It is easy to show that,

$$\mathbb{E}_q[\log q(\boldsymbol{\Theta})] = \sum_{j=1}^{p-1}\sum_{t=1}^{k}\left(\mathbb{E}_q[\log q(\beta_{jt})] + \mathbb{E}_q[\log q(\delta_{jt})]\right) + \sum_{j=1}^{p-1}\left(\mathbb{E}_q[\log q(\kappa_j)] + \mathbb{E}_q[\log q(\beta_{0j})]\right) + \sum_{i=1}^{n}\sum_{t=1}^{k}\mathbb{E}_q[\log q(F_{it})] + \sum_{i=1}^{n}\sum_{j=1}^{p-1}\mathbb{E}_q[\log q(z_{ij})]. \quad (1)$$

Equation (1) further implies that,

$$\begin{aligned}\mathbb{E}_q[\log q(\boldsymbol{\Theta})] &= \sum_{j=1}^{p-1}\sum_{t=1}^{k}\Big(-\frac{1}{2}\log\lambda_{jt}^2 - g_{1jt} + \log g_{2jt} - \log\mathit{\Gamma} g_{1jt} - (1-g_{1jt})\psi(g_{1jt})\Big)\\ &+ \sum_{j=1}^{p-1}\Big((\tau_{j1}-1)\{\psi(\tau_{j1})-\psi(\tau_{j1}+\tau_{j2})\} + (\tau_{j2}-1)\{\psi(\tau_{j2})-\psi(\tau_{j1}+\tau_{j2})\}\\ &- \log\mathcal{B}(\tau_{j1},\tau_{j2})\Big) - \frac{1}{2}\sum_{j=1}^{p-1}\log C_{0j}^2 - \sum_{i=1}^{n}\sum_{t=1}^{k}\left(\log\omega_{it} - \mathit{\Psi}_2(\alpha_{it})\right)\\ &+ \sum_{i=1}^{n}\sum_{j=1}^{p-1}\left(\pi_{ij}\log\pi_{ij} + (1-\pi_{ij})\log(1-\pi_{ij})\right)\end{aligned}$$

where,

$$\begin{aligned}\mathit{\Psi}_2(\alpha_{it}) &= \mathbb{E}_q\left(\log\left(\mathit{\Phi}\left[\alpha_{it}\left(\frac{F_{it}-\xi_{it}}{\omega_{it}}\right)\right]\right)\right)\\ &= \int_{\mathbb{R}}\frac{2}{\omega_{it}}\phi\left(\frac{f_{it}-\xi_{it}}{\omega_{it}}\right)\mathit{\Phi}\left[\alpha_{it}\left(\frac{f_{it}-\xi_{it}}{\omega_{it}}\right)\right]\log\left(\mathit{\Phi}\left[\alpha_{it}\left(\frac{f_{it}-\xi_{it}}{\omega_{it}}\right)\right]\right)df_{it}\end{aligned}$$

Substituting $\frac{f_{it}-\xi_{it}}{\omega_{it}} = u_{it}$, we get,

$$\mathit{\Psi}_2(\alpha_{it}) = \int_{\mathbb{R}} 2\phi(u_{it})\mathit{\Phi}(\alpha_{it}u_{it})\log\mathit{\Phi}(\alpha_{it}u_{it})du_{it} = \mathbb{E}[\log\mathit{\Phi}(\alpha_{it}U_{it})]$$

with $U_{it} \sim \mathcal{SN}(0,1,\alpha_{it})$.

Since, the closed form for $\mathit{\Psi}_2(\alpha_{it})$ does not exist, we use a Taylor expansion, and denote $\mathit{\Psi}_2(\alpha_{it}) \approx \mathit{\Psi}_2^{AP}(\alpha_{it})$. The details can be found in section 6.

#### 1.2 Derivation of $\mathbb{E}_q[\log p(\boldsymbol{x}, \boldsymbol{\Theta})]$

$$
\begin{aligned}
\mathbb{E}_q[\log p(\boldsymbol{x}, \boldsymbol{\Theta})] &= \sum_{i=1}^{n} \mathbb{E}_q[\log p(\boldsymbol{x}_i \mid \boldsymbol{\rho}_i, M_i)] + \sum_{j=1}^{p-1}\sum_{t=1}^{k} \left(\mathbb{E}_q[\log p(\beta_{jt} \mid \delta_{jt})] + \mathbb{E}_q[\log p(\delta_{jt})]\right) \\
&+ \sum_{j=1}^{p-1} \left(\mathbb{E}_q[\log p(\kappa_j)] + \mathbb{E}_q[\log p(\beta_{0j})]\right) + \sum_{i=1}^{n}\sum_{t=1}^{k} \mathbb{E}_q[\log p(F_{it})] \\
&+ \sum_{i=1}^{n}\sum_{j=1}^{p-1} \mathbb{E}_q[\log p(z_{ij} \mid \kappa_j)]
\end{aligned}
$$

where,

$$
\begin{aligned}
\mathbb{E}_q[\log p(\beta_{jt} \mid \delta_{jt})] &= \frac{1}{2}\left(\psi(g_{1jt}) - \log g_{2jt}\right) - \frac{1}{2}\left(\lambda_{jt}^2 + r_{jt}^2\right)\frac{g_{1jt}}{g_{2jt}} \\
\mathbb{E}_q[\log p(\delta_{jt})] &= (g_1 - 1)\left(\psi(g_{1jt}) - \log g_{2jt}\right) - g_2\frac{g_{1jt}}{g_{2jt}} \\
\mathbb{E}_q[\log p(\kappa_j)] &= (\nu_1 - 1)\{\psi(\tau_{j1}) - \psi(\tau_{j1} + \tau_{j2})\} + (\nu_2 - 1)\{\psi(\tau_{j2}) - \psi(\tau_{j1} + \tau_{j2})\} \\
\mathbb{E}_q[\log p(\beta_{0j})] &= -\frac{1}{2}\left(a_{0j}^2 + C_{0j}^2\right) \\
\mathbb{E}_q[\log p(F_{it})] &= -\frac{1}{2}\left(\omega_{it}^2 + \xi_{it}^2 + 2\sqrt{\frac{2}{\pi}}\frac{\xi_{it}\omega_{it}\alpha_{it}}{\sqrt{1+\alpha_{it}^2}} - 2\alpha^*\left(\xi_{it} + \frac{\omega_{it}\alpha_{it}}{\sqrt{1+\alpha_{it}^2}}\sqrt{\frac{2}{\pi}}\right)\right) \\
&+ \varPsi_1(\xi_{it}, \omega_{it}, \alpha_{it}) \\
\mathbb{E}_q[\log p(z_{ij} \mid \kappa_j)] &= \pi_{ij}\{\psi(\tau_{j1}) - \psi(\tau_{j1} + \tau_{j2})\} + (1 - \pi_{ij})\{\psi(\tau_{j2}) - \psi(\tau_{j1} + \tau_{j2})\}
\end{aligned}
$$

Note that, $\varPsi_1(\xi_{it}, \omega_{it}, \alpha_{it}) = \mathbb{E}_q[\log \varPhi(\alpha_0(F_{it} - \alpha^*))]$. Again, since, the closed form for $\varPsi_1(\xi_{it}, \omega_{it}, \alpha_{it})$ does not exist, we use a Taylor expansion, and denote $\varPsi_1(\xi_{it}, \omega_{it}, \alpha_{it}) \approx \varPsi_1^{AP}(\xi_{it}, \omega_{it}, \alpha_{it})$. The details can be found in section 5.

Now, let's find $\mathbb{E}_q[\log p(\boldsymbol{x}_i \mid \boldsymbol{\rho}_i, M_i)]$.

$$
\log \rho_{ij} = \log(1 - z_{ij}) + \beta_{0j} + \boldsymbol{F}_i^T\boldsymbol{\beta}_j - \log\left(1 + \sum_{\ell=1}^{p-1}(1 - z_{i\ell})\exp\left(\beta_{0\ell} + \boldsymbol{F}_i^T\boldsymbol{\beta}_\ell\right)\right),
$$

$$
x_{ij}\log \rho_{ij} = x_{ij}\log(1 - z_{ij}) + x_{ij}\left(\beta_{0j} + \sum_{t=1}^{k} F_{it}\beta_{jt}\right) - x_{ij}\log\left(1 + \sum_{\ell=1}^{p-1}(1 - z_{i\ell})\exp\left(\beta_{0\ell} + \boldsymbol{F}_i^T\boldsymbol{\beta}_\ell\right)\right).
$$

Since, $x_{ij}\log(1-z_{ij})=0$,

$$\begin{aligned}
\sum_{j=1}^{p}\mathbb{E}_q[x_{ij}\log\rho_{ij}] &= \sum_{j=1}^{p} x_{ij}\left(\mathbb{E}_q[\beta_{0j}]+\sum_{t=1}^{k}\mathbb{E}_q[F_{it}]\mathbb{E}_q[\beta_{jt}]\right)\\
&-\sum_{j=1}^{p} x_{ij}\mathbb{E}_q\left[\log\left(1+\sum_{\ell=1}^{p-1}(1-z_{i\ell})\exp\left(\beta_{0\ell}+\boldsymbol{F}_i^T\boldsymbol{\beta}_\ell\right)\right)\right]\\
&\geq \sum_{j=1}^{p-1} x_{ij}\left[a_{0j}+\sum_{t=1}^{k}\left(\xi_{it}+\frac{\omega_{it}\alpha_{it}}{\sqrt{1+\alpha_{it}^2}}\sqrt{\frac{2}{\pi}}\right)(r_{jt})\right]\\
&- M_i\log\left(1+\sum_{\ell=1}^{p-1}(1-\pi_{i\ell})\mathbb{E}_q[\exp(\beta_{0\ell})]\prod_{t=1}^{k}\mathbb{E}_q\left[\exp\left(F_{it}\beta_{\ell t}\right)\right]\right)\\
&= \sum_{j=1}^{p-1} x_{ij}\left[a_{0j}+\sum_{t=1}^{k}\left(\xi_{it}+\frac{\omega_{it}\alpha_{it}}{\sqrt{1+\alpha_{it}^2}}\sqrt{\frac{2}{\pi}}\right)(r_{jt})\right]\\
&- M_i\log\left(1+\sum_{\ell=1}^{p-1}(1-\pi_{i\ell})\exp\left(a_{0\ell}+\frac{1}{2}C_{0\ell}^2+\boldsymbol{L}_{i\ell}\right)\right)
\end{aligned}$$

where,

$$\begin{aligned}
\boldsymbol{L}_{i\ell} &:= \sum_{t=1}^{k}\log L_{i\ell t}\\
&= \sum_{t=1}^{k}\log\left(\frac{2}{\sqrt{1-\lambda_{\ell t}^2\omega_{it}^2}}\times\exp\left\{\frac{\lambda_{\ell t}^2\xi_{it}^2+r_{\ell t}^2\omega_{it}^2+2r_{\ell t}\xi_{it}}{2(1-\lambda_{\ell t}^2\omega_{it}^2)}\right\}\times\Phi\left[\frac{(r_{\ell t}+\lambda_{\ell t}^2\xi_{it})\omega_{it}\alpha_{it}}{\sqrt{1-\lambda_{\ell t}^2\omega_{it}^2}\sqrt{1+\alpha_{it}^2-\lambda_{\ell t}^2\omega_{it}^2}}\right]\right).
\end{aligned}$$

Therefore,

$$\begin{aligned}
\mathbb{E}_q[\log p(\boldsymbol{x}_i\mid\boldsymbol{\rho}_i,M_i)] &= \sum_{j=1}^{p}\mathbb{E}_q[x_{ij}\log\rho_{ij}]\\
&\geq \sum_{j=1}^{p-1} x_{ij}\left[a_{0j}+\sum_{t=1}^{k}\left(\xi_{it}+\frac{\omega_{it}\alpha_{it}}{\sqrt{1+\alpha_{it}^2}}\sqrt{\frac{2}{\pi}}\right)(r_{jt})\right]\\
&- M_i\log\left(1+\sum_{\ell=1}^{p-1}(1-\pi_{i\ell})\exp\left(a_{0\ell}+\frac{1}{2}C_{0\ell}^2+\boldsymbol{L}_{i\ell}\right)\right)
\end{aligned}$$

Hence, final expression for $\mathbb{E}_q[\log p(\boldsymbol{x}, \boldsymbol{\Theta})]$ is,

$$
\begin{aligned}
\mathbb{E}_q[\log p(\boldsymbol{x}, \boldsymbol{\Theta})] \approx & \sum_{i=1}^{n}\sum_{j=1}^{p-1} x_{ij} a_{0j} + \sum_{i=1}^{n}\sum_{j=1}^{p-1}\sum_{t=1}^{k} x_{ij} r_{jt} \left( \xi_{it} + \frac{\omega_{it}\alpha_{it}}{\sqrt{1+\alpha_{it}^2}} \sqrt{\frac{2}{\pi}} \right) \\
& - \sum_{i=1}^{n} M_i \log \left( 1 + \sum_{\ell=1}^{p-1} (1 - \pi_{i\ell}) \exp \left( a_{0\ell} + \frac{1}{2} C_{0\ell}^2 + \boldsymbol{L}_{i\ell} \right) \right) \\
& + \sum_{j=1}^{p-1}\sum_{t=1}^{k} \left( \left( g_1 - \frac{1}{2} \right) (\psi(g_{1jt}) - \log g_{2jt}) - g_2 \frac{g_{1jt}}{g_{2jt}} - \frac{1}{2} \left( \lambda_{jt}^2 + r_{jt}^2 \right) \frac{g_{1jt}}{g_{2jt}} \right) \\
& - \sum_{i=1}^{n}\sum_{t=1}^{k} \left( \frac{\omega_{it}^2}{2} + \frac{\xi_{it}^2}{2} + \sqrt{\frac{2}{\pi}} \frac{\xi_{it}\omega_{it}\alpha_{it}}{\sqrt{1+\alpha_{it}^2}} - \alpha^* \left( \xi_{it} + \frac{\omega_{it}\alpha_{it}}{\sqrt{1+\alpha_{it}^2}} \sqrt{\frac{2}{\pi}} \right) \right. \\
& \left. - \Psi_1^{AP}(\xi_{it}, \omega_{it}, \alpha_{it}) \right) + \sum_{j=1}^{p-1} \Bigg( (\nu_1 - 1)\{\psi(\tau_{j1}) - \psi(\tau_{j1} + \tau_{j2})\} \\
& + (\nu_2 - 1)\{\psi(\tau_{j2}) - \psi(\tau_{j1} + \tau_{j2})\} - \frac{1}{2} \left( a_{0j}^2 + C_{0j}^2 \right) \Bigg) \\
& + \sum_{i=1}^{n}\sum_{j=1}^{p-1} \left[ \pi_{ij}\{\psi(\tau_{j1}) - \psi(\tau_{j1} + \tau_{j2})\} + (1 - \pi_{ij})\{\psi(\tau_{j2}) - \psi(\tau_{j1} + \tau_{j2})\} \right].
\end{aligned}
$$

Finally,

$$
\mathrm{ELBO} = \mathbb{E}_q[\log p(\boldsymbol{x}, \boldsymbol{\Theta}) - \log q(\boldsymbol{\Theta})] = \mathbb{E}_q[\log p(\boldsymbol{x}, \boldsymbol{\Theta})] - \mathbb{E}_q[\log q(\boldsymbol{\Theta})] \approx \mathrm{ELBO}^{\mathrm{AP}}
$$

#### 2 MGF of Product of Normal and Skew Normal Random Variables

**Lemma 1.** *Let $U \sim \mathcal{N}(0,1)$, and $h,k \in \mathbb{R}$. Then,*

$$\mathbb{E}\left[\Phi\left(hU+k\right)\right] = \Phi\left[\frac{k}{\sqrt{1+h^2}}\right]$$

*where, $\Phi(.)$ is the standard normal cdf.*

*Proof.* $\forall\ h,k \in \mathbb{R}$, we denote $\mathbb{E}\left[\Phi\left(hU+k\right)\right]$ as,

$$\Psi(h,k) = \mathbb{E}\left[\Phi\left(hU+k\right)\right] = \int_{\mathbb{R}} \Phi\left(hu+k\right)\phi(u)du \tag{2}$$

where $\phi(.)$ is the standard normal pdf.

Differentiating equation (2) with respect to $k$ we get,

$$\begin{aligned}
\frac{\partial \Psi(h,k)}{\partial k} &= \int_{\mathbb{R}} \phi\left(hu+k\right)\phi(u)du \\
&= \frac{1}{2\pi}\int_{\mathbb{R}} \exp\left(-\frac{1}{2}\left[(hu+k)^2+u^2\right]\right)du \\
&= \frac{1}{2\pi}\exp\left(-\frac{k^2}{2(1+h^2)}\right)\int_{\mathbb{R}} \exp\left(-\frac{(1+h^2)}{2}\left(u+\frac{hk}{1+h^2}\right)^2\right)du
\end{aligned}$$

Letting $v = \sqrt{1+h^2}\left(u+\frac{hk}{1+h^2}\right)$ we have,

$$\begin{aligned}
\frac{\partial \Psi(h,k)}{\partial k} &= \frac{1}{\sqrt{2\pi}\sqrt{1+h^2}}\exp\left(-\frac{k^2}{2(1+h^2)}\right)\frac{1}{\sqrt{2\pi}}\int_{\mathbb{R}}\exp\left(-\frac{v^2}{2}\right)dv \\
&= \frac{1}{\sqrt{1+h^2}}\phi\left(\frac{k}{\sqrt{1+h^2}}\right)
\end{aligned}$$

Now integrating with respect to $k$ we get,

$$\Psi(h,k) = \mathbb{E}\left[\Phi\left(hU+k\right)\right] = \Phi\left[\frac{k}{\sqrt{1+h^2}}\right]$$

□

**Lemma 2.** *Let $X \sim \mathcal{N}(\mu,\sigma^2)$, $Y \sim \mathcal{SN}(\xi,\omega,\alpha)$, X & Y are independent, and $Z = XY$. If $|s| < \frac{1}{\sigma\omega}$, then the MGF of Z is given by,*

$$M_Z(s) = \frac{2}{\sqrt{1-\sigma^2\omega^2 s^2}} \times \exp\left(\frac{(\sigma^2\xi^2+\mu^2\omega^2)s^2+2\mu\xi s}{2(1-\sigma^2\omega^2 s^2)}\right) \times \Phi\left[\frac{(\mu+\sigma^2\xi s)\omega\delta s}{\sqrt{1-\sigma^2\omega^2 s^2}\sqrt{1-\sigma^2\omega^2 s^2(1-\delta^2)}}\right]$$

*where, $\Phi$ is the CDF of the standard normal distribution and $\delta = \frac{\alpha}{\sqrt{1+\alpha^2}}$.*

*Proof.*

$$M_Z(s) = M_{XY}(s) = \mathbb{E}\left[e^{sXY}\right]$$

Using law of total expectation,

$$\mathbb{E}\left[e^{sXY}\right] = \mathbb{E}\left[\mathbb{E}\left[e^{sXY} \mid X\right]\right]$$

Finding the inner expectation first, we have,

$$\mathbb{E}[e^{sXY} \mid X] = 2\exp\left(\xi sx + \frac{\omega^2 s^2 x^2}{2}\right)\Phi(\omega\delta sx) \tag{3}$$

where equation (2) is the MGF of skew-normal random variable. Hence,

$$\begin{aligned}
\mathbb{E}[e^{sXY}] &= \mathbb{E}\left[2\exp\left(\xi sX + \frac{\omega^2 s^2 X^2}{2}\right)\Phi(\omega\delta sX)\right] \\
&= \frac{2}{\sqrt{2\pi}\sigma}\int_{-\infty}^{\infty}\exp\left(\xi sx + \frac{\omega^2 s^2 x^2}{2} - \frac{1}{2}\left(\frac{x-\mu}{\sigma}\right)^2\right)\Phi(\omega\delta sx)dx \\
&= \frac{2\exp\left(-\frac{\mu^2}{2\sigma^2} + \frac{B^2}{2A}\right)}{\sqrt{2\pi}\sigma}\int_{-\infty}^{\infty}\exp\left(-\frac{A}{2}\left(x - \frac{B}{A}\right)^2\right)\Phi(Dx)dx
\end{aligned}$$

where, $A = \frac{1}{\sigma^2} - \omega^2 s^2$, $B = \frac{\mu}{\sigma^2} + \xi s$, and $D = \omega\delta s$.

Now, substituting $\sqrt{A}\left(x - \frac{B}{A}\right) = u$, we have,

$$\begin{aligned}
\mathbb{E}[e^{sXY}] &= \frac{2\exp\left(-\frac{\mu^2}{2\sigma^2} + \frac{B^2}{2A}\right)}{\sqrt{2\pi}\sigma\sqrt{A}}\int_{-\infty}^{\infty}\exp\left(-\frac{u^2}{2}\right)\Phi\left[D\left(\frac{u}{\sqrt{A}} + \frac{B}{A}\right)\right]du \\
&= \frac{2\exp\left(-\frac{\mu^2}{2\sigma^2} + \frac{B^2}{2A}\right)}{\sigma\sqrt{A}}\mathbb{E}\left[\Phi\left(U\frac{D}{\sqrt{A}} + \frac{BD}{A}\right)\right]
\end{aligned}$$

where $U \sim \mathcal{N}(0,1)$.

Therefore, using Lemma 1, we have,

$$\mathbb{E}[e^{sXY}] = \frac{2\exp\left(-\frac{\mu^2}{2\sigma^2} + \frac{B^2}{2A}\right)}{\sigma\sqrt{A}}\Phi\left[\frac{\frac{BD}{A}}{\sqrt{1 + \frac{D^2}{A}}}\right]$$

$$\mathbb{E}[e^{sXY}] = \frac{2}{\sqrt{1 - \sigma^2\omega^2 s^2}} \times \exp\left(\frac{(\sigma^2\xi^2 + \mu^2\omega^2)s^2 + 2\mu\xi s}{2(1 - \sigma^2\omega^2 s^2)}\right) \times \Phi\left[\frac{(\mu + \sigma^2\xi s)\omega\delta s}{\sqrt{1 - \sigma^2\omega^2 s^2}\sqrt{1 - \sigma^2\omega^2 s^2(1 - \delta^2)}}\right]$$

□

### 3 Parameter Updates

Denote the matrices as, $\boldsymbol{R} = (r_{jt}) \in \mathbb{R}^{(p-1)\times k}$, $\boldsymbol{\Lambda} = (\lambda_{jt}^2) \in (0,1)^{(p-1)\times k}$, $\boldsymbol{G}_1 = (g_{1jt}) \in \mathbb{R}_+^{(p-1)\times k}$, $\boldsymbol{G}_2 = (g_{2jt}) \in \mathbb{R}_+^{(p-1)\times k}$ where $\mathbb{R}_+$ is the set of all positive reals, $\boldsymbol{\Xi} = (\xi_{it}) \in \mathbb{R}^{n\times k}$, $\boldsymbol{\Omega} = (\omega_{it}) \in (0,1)^{n\times k}$, and $\boldsymbol{A} = (\alpha_{it}) \in \mathbb{R}^{n\times k}$. Let the vectors be denoted as, $\boldsymbol{A}_0 = (a_{01}, \dots, a_{0(p-1)})^T \in \mathbb{R}^{(p-1)}$ and $\boldsymbol{C}_0 = (C_{01}^2, \dots, C_{0(p-1)}^2)^T \in \mathbb{R}_+^{(p-1)}$. Moreover, $\odot$ is Hadamard (elementwise) product, $\oslash$ is Hadamard division, and usual matrix multiplication will be denoted as $\boldsymbol{XY}$ where $\boldsymbol{X}$ and $\boldsymbol{Y}$ are compatible matrices.

Now, We formulate the optimization for all variational parameters (except $\pi_{ij}$) by defining the corresponding objective and gradient functions, performing estimation via the `optim` function in R. While the standard BFGS quasi-Newton algorithm is employed for unconstrained variational parameters, the L-BFGS-B method is utilized where bounds are required. Specifically, we enforce constraints on $\boldsymbol{G}_1$, $\boldsymbol{G}_2$, $\boldsymbol{C}_0$, and $\{\tau_{j1}, \tau_{j2}\}_{j=1}^{p-1}$ to satisfy the positivity requirements of their respective variational Beta and Gamma distributions. Additionally, the elements of $\boldsymbol{\Lambda}$ and $\boldsymbol{\Omega}$ are restricted to the $(0,1)$ to ensure well defined $\boldsymbol{L}_{ij}$.

1. Update $\tau_{j1}^{(s)}$

$$\tau_{j1}^{(s)} = \underset{\tau_{j1}}{\arg\max}\ \mathrm{ELBO}^{\mathrm{AP}}(\tau_{j1})$$

with objective function,

$$\begin{aligned}\mathrm{ELBO}^{\mathrm{AP}}(\tau_{j1}) = &\sum_{i=1}^{n}\left\{\pi_{ij}^{(s)}\psi(\tau_{j1}) - \psi\left(\tau_{j1}+\tau_{j2}^{(s-1)}\right)\right\} + (\nu_1-\tau_{j1})\psi(\tau_{j1}) \\ &+ \psi\left(\tau_{j1}+\tau_{j2}^{(s-1)}\right)\left(\tau_{j1}-\nu_1+\tau_{j2}^{(s-1)}-\nu_2\right) + \log\mathcal{B}\left(\tau_{j1},\tau_{j2}^{(s-1)}\right)\end{aligned}$$

and gradient function,

$$\begin{aligned}\frac{\partial}{\partial\tau_{j1}}\mathrm{ELBO}^{\mathrm{AP}}(\tau_{j1}) = &\sum_{i=1}^{n}\left\{\pi_{ij}^{(s)}\psi^{(1)}(\tau_{j1}) - \psi^{(1)}\left(\tau_{j1}+\tau_{j2}^{(s-1)}\right)\right\} + (\nu_1-\tau_{j1})\psi^{(1)}(\tau_{j1}) \\ &+ \psi^{(1)}\left(\tau_{j1}+\tau_{j2}^{(s-1)}\right)\left(\tau_{j1}-\nu_1+\tau_{j2}^{(s-1)}-\nu_2\right)\end{aligned}$$

2. Update $\tau_{j2}^{(s)}$

$$\tau_{j2}^{(s)} = \underset{\tau_{j2}}{\arg\max}\ \mathrm{ELBO}^{\mathrm{AP}}(\tau_{j2})$$

with objective function,

$$\begin{aligned}\mathrm{ELBO}^{\mathrm{AP}}(\tau_{j2}) = &\sum_{i=1}^{n}\left\{\left(1-\pi_{ij}^{(s)}\right)\psi(\tau_{j2}) - \psi\left(\tau_{j1}^{(s)}+\tau_{j2}\right)\right\} + (\nu_2-\tau_{j2})\psi(\tau_{j2}) \\ &+ \psi\left(\tau_{j1}^{(s)}+\tau_{j2}\right)\left(\tau_{j1}^{(s)}-\nu_1+\tau_{j2}-\nu_2\right) + \log\mathcal{B}\left(\tau_{j1}^{(s)},\tau_{j2}\right)\end{aligned}$$

and gradient function,

$$\begin{aligned}\frac{\partial}{\partial\tau_{j2}}\mathrm{ELBO}^{\mathrm{AP}}(\tau_{j2}) = &\sum_{i=1}^{n}\left\{\left(1-\pi_{ij}^{(s)}\right)\psi^{(1)}(\tau_{j2}) - \psi^{(1)}\left(\tau_{j1}^{(s)}+\tau_{j2}\right)\right\} + (\nu_2-\tau_{j2})\psi^{(1)}(\tau_{j2}) \\ &+ \psi^{(1)}\left(\tau_{j1}^{(s)}+\tau_{j2}\right)\left(\tau_{j1}^{(s)}-\nu_1+\tau_{j2}-\nu_2\right)\end{aligned}$$

3. Update $\boldsymbol{G}_1^{(s)}$

$$\boldsymbol{G}_1^{(s)} = \underset{\boldsymbol{G}_1}{\arg\max}\ \text{ELBO}^{\text{AP}}(\boldsymbol{G}_1)$$

with objective function,

$$\begin{aligned}\text{ELBO}^{\text{AP}}(\boldsymbol{G}_1) = &\sum_{j=1}^{p-1}\sum_{t=1}^{k}\left(g_1 - \frac{1}{2}\right)\psi(g_{1jt}) - \frac{g_{1jt}}{g_{2jt}^{(s-1)}}\left(\frac{1}{2}\left(\left(\lambda_{jt}^2\right)^{(s-1)} + \left(r_{jt}^2\right)^{(s-1)}\right) + g_2\right)\\ &+ g_{1jt} + \log \mathit{\Gamma} g_{1jt} + (1 - g_{1jt})\psi(g_{1jt})\end{aligned}$$

and gradient function,

$$\begin{aligned}\frac{\partial}{\partial \boldsymbol{G}_1}\text{ELBO}^{\text{AP}}(\boldsymbol{G}_1) = &\left(g_1 - \frac{1}{2}\right)\psi^{(1)}(\boldsymbol{G}_1) - \frac{1}{\boldsymbol{G}_2^{(s-1)}}\left(\frac{1}{2}\left(\boldsymbol{\Lambda}^{(s-1)} + (\boldsymbol{R}\odot\boldsymbol{R})^{(s-1)}\right) + g_2\right) + 1\\ &+ (1 - \boldsymbol{G}_1)\psi^{(1)}(\boldsymbol{G}_1)\end{aligned}$$

4. Update $\boldsymbol{G}_2^{(s)}$

$$\boldsymbol{G}_2^{(s)} = \underset{\boldsymbol{G}_2}{\arg\max}\ \text{ELBO}^{\text{AP}}(\boldsymbol{G}_2)$$

with objective function,

$$\begin{aligned}\text{ELBO}^{\text{AP}}(\boldsymbol{G}_2) = &-\sum_{j=1}^{p-1}\sum_{t=1}^{k}\left(g_1 - \frac{1}{2}\right)\log g_{2jt} - \frac{g_{1jt}^{(s)}}{g_{2jt}}\left(\frac{1}{2}\left(\left(\lambda_{jt}^2\right)^{(s-1)} + \left(r_{jt}^2\right)^{(s-1)}\right) + g_2\right)\\ &- \log(g_{2jt})\end{aligned}$$

and gradient function,

$$\frac{\partial}{\partial \boldsymbol{G}_2}\text{ELBO}^{\text{AP}}(\boldsymbol{G}_2) = -\left(g_1 + \frac{1}{2}\right)\frac{1}{\boldsymbol{G}_2} + \frac{\boldsymbol{G}_1^{(s)}}{\boldsymbol{G}_2^2}\left(\frac{1}{2}\left(\boldsymbol{\Lambda}^{(s-1)} + (\boldsymbol{R}\odot\boldsymbol{R})^{(s-1)}\right) + g_2\right)$$

5. Update $\boldsymbol{R}^{(s)}$

$$\boldsymbol{R}^{(s)} = \underset{\boldsymbol{R}}{\arg\max}\ \text{ELBO}^{\text{AP}}(\boldsymbol{R})$$

with objective function,

$$\begin{aligned}\text{ELBO}^{\text{AP}}(\boldsymbol{R}) = &\sum_{i=1}^{n}\sum_{j=1}^{p-1}\sum_{t=1}^{k} x_{ij} r_{jt}\left(\xi_{it}^{(s-1)} + \frac{\omega_{it}^{(s-1)}\alpha_{it}^{(s-1)}}{\sqrt{1 + \left(\alpha_{it}^2\right)^{(s-1)}}}\sqrt{\frac{2}{\pi}}\right)\\ &- \sum_{i=1}^{n} M_i \log\left(1 + \sum_{\ell=1}^{p-1}\left(1 - \pi_{i\ell}^{(s)}\right)\exp\left(a_{0\ell}^{(s-1)} + \frac{1}{2}\left(C_{0\ell}^2\right)^{(s-1)} + \boldsymbol{L}_{i\ell}^{(s-1)}\right)\right)\\ &- \frac{1}{2}\sum_{j=1}^{p-1}\sum_{t=1}^{k} r_{jt}^2\left(\frac{g_{1jt}^{(s)}}{g_{2jt}^{(s)}}\right)\end{aligned}$$

and gradient function,

$$\frac{\partial}{\partial \boldsymbol{R}}\text{ELBO}^{\text{AP}}(\boldsymbol{R}) = X^T \left( \boldsymbol{\Xi}^{(s-1)} + \sqrt{\frac{2}{\pi}} \left( \boldsymbol{\Omega}^{(s-1)} \odot \boldsymbol{A}^{(s-1)} \right) \oslash \sqrt{1 + (\boldsymbol{A} \odot \boldsymbol{A})^{(s-1)}} \right) - \boldsymbol{R} \odot \left( \boldsymbol{G}_1^{(s)} \oslash \boldsymbol{G}_2^{(s)} \right) - \widetilde{\boldsymbol{R}}$$

where, the elements of matrix $\widetilde{\boldsymbol{R}}$,

$$\widetilde{R}_{jt} = \sum_{i=1}^{n} M_i \left( \frac{\left(1 - \pi_{ij}^{(s)}\right) H_{ijt}^{(s-1)} \exp\left( a_{0j}^{(s-1)} + \frac{1}{2} \left( C_{0j}^2 \right)^{(s-1)} + \boldsymbol{L}_{ij}^{(s-1)} \right)}{1 + \sum_{\ell=1}^{p-1} \left(1 - \pi_{i\ell}^{(s)}\right) \exp\left( a_{0\ell}^{(s-1)} + \frac{1}{2} \left( C_{0\ell}^2 \right)^{(s-1)} + \boldsymbol{L}_{i\ell}^{(s-1)} \right)} \right)$$

and,

$$H_{ijt}^{(s-1)} = \frac{\partial}{\partial r_{jt}} \boldsymbol{L}_{ij}^{(s-1)}$$

6. Update $\boldsymbol{\Lambda}^{(s)}$

$$\boldsymbol{\Lambda}^{(s)} = \arg\max_{\boldsymbol{\Lambda}} \text{ELBO}^{\text{AP}}(\boldsymbol{\Lambda})$$

with objective function,

$$\text{ELBO}^{\text{AP}}(\boldsymbol{\Lambda}) = -\sum_{i=1}^{n} M_i \log \left( 1 + \sum_{\ell=1}^{p-1} \left(1 - \pi_{i\ell}^{(s)}\right) \exp\left( a_{0\ell}^{(s-1)} + \frac{1}{2} \left( C_{0\ell}^2 \right)^{(s-1)} + \boldsymbol{L}_{i\ell}^{(s-1)} \right) \right) - \frac{1}{2} \sum_{j=1}^{p-1} \sum_{t=1}^{k} \left( \lambda_{jt}^2 \left( \frac{g_{1jt}^{(s)}}{g_{2jt}^{(s)}} \right) - \log \lambda_{jt}^2 \right)$$

and gradient function,

$$\frac{\partial}{\partial \boldsymbol{\Lambda}}\text{ELBO}^{\text{AP}}(\boldsymbol{\Lambda}) = -\widetilde{\boldsymbol{N}} - \frac{1}{2} \left( \boldsymbol{G}_1^{(s)} \oslash \boldsymbol{G}_2^{(s)} - 1 \oslash \boldsymbol{\Lambda} \right)$$

where, the elements of matrix $\widetilde{\boldsymbol{N}}$,

$$\widetilde{N}_{jt} = \sum_{i=1}^{n} M_i \left( \frac{\left(1 - \pi_{ij}^{(s)}\right) M_{ijt}^{(s-1)} \exp\left( a_{0j}^{(s-1)} + \frac{1}{2} \left( C_{0j}^2 \right)^{(s-1)} + \boldsymbol{L}_{ij}^{(s-1)} \right)}{1 + \sum_{\ell=1}^{p-1} \left(1 - \pi_{i\ell}^{(s)}\right) \exp\left( a_{0\ell}^{(s-1)} + \frac{1}{2} \left( C_{0\ell}^2 \right)^{(s-1)} + \boldsymbol{L}_{i\ell}^{(s-1)} \right)} \right)$$

and,

$$M_{ijt}^{(s-1)} = \frac{\partial}{\partial \lambda_{jt}^2} \boldsymbol{L}_{ij}^{(s-1)}$$

7. Update $\boldsymbol{\Xi}^{(s)}$

$$\boldsymbol{\Xi}^{(s)} = \arg\max_{\boldsymbol{\Xi}} \text{ELBO}^{\text{AP}}(\boldsymbol{\Xi})$$

with objective function,

$$\begin{aligned}
\text{ELBO}^{\text{AP}}(\boldsymbol{\Xi}) &= \sum_{i=1}^{n}\sum_{j=1}^{p-1}\sum_{t=1}^{k} x_{ij}\xi_{it}r_{jt}^{(s)} \\
&\quad - \sum_{i=1}^{n} M_i \log\left(1 + \sum_{\ell=1}^{p-1}\left(1-\pi_{i\ell}^{(s)}\right)\exp\left(a_{0\ell}^{(s-1)} + \frac{1}{2}\left(C_{0\ell}^2\right)^{(s-1)} + \boldsymbol{L}_{i\ell}^{(s-1)}\right)\right) \\
&\quad - \sum_{i=1}^{n}\sum_{t=1}^{k}\left(\frac{\xi_{it}^2}{2} + \sqrt{\frac{2}{\pi}}\frac{\xi_{it}\omega_{it}^{(s-1)}\alpha_{it}^{(s-1)}}{\sqrt{1+(\alpha_{it}^2)^{(s-1)}}} - \alpha^*\xi_{it} - \Psi_1^{AP}\left(\xi_{it}, \omega_{it}^{(s-1)}, \alpha_{it}^{(s-1)}\right)\right)
\end{aligned}$$

and gradient function,

$$\begin{aligned}
\frac{\partial}{\partial\boldsymbol{\Xi}}\text{ELBO}^{\text{AP}}(\boldsymbol{\Xi}) &= X\boldsymbol{R}^{(s)} - \widetilde{\boldsymbol{P}} - \left(\boldsymbol{\Xi} + \sqrt{\frac{2}{\pi}}\left(\boldsymbol{\Omega}^{(s-1)} \odot \boldsymbol{A}^{(s-1)}\right) \oslash \sqrt{1 + (\boldsymbol{A}\odot\boldsymbol{A})^{(s-1)}}\right) \\
&\quad + \alpha^* + \frac{\partial}{\partial\boldsymbol{\Xi}}\boldsymbol{\Psi}_1^{AP}\left(\boldsymbol{\Xi}, \boldsymbol{\Omega}^{(s-1)}, \boldsymbol{A}^{(s-1)}\right)
\end{aligned}$$

where, the elements of matrix $\widetilde{\boldsymbol{P}}$,

$$\widetilde{P}_{it} = M_i\left(\frac{\sum_{j=1}^{p-1}\left(1-\pi_{ij}^{(s)}\right)E_{ijt}^{(s-1)}\exp\left(a_{0j}^{(s-1)} + \frac{1}{2}\left(C_{0j}^2\right)^{(s-1)} + \boldsymbol{L}_{ij}^{(s-1)}\right)}{1 + \sum_{\ell=1}^{p-1}\left(1-\pi_{i\ell}^{(s)}\right)\exp\left(a_{0\ell}^{(s-1)} + \frac{1}{2}\left(C_{0\ell}^2\right)^{(s-1)} + \boldsymbol{L}_{i\ell}^{(s-1)}\right)}\right)$$

and,

$$E_{ijt}^{(s-1)} = \frac{\partial}{\partial\xi_{it}}\boldsymbol{L}_{ij}^{(s-1)}$$

8. Update $\boldsymbol{\Omega}^{(s)}$

$$\boldsymbol{\Omega}^{(s)} = \arg\max_{\boldsymbol{\Omega}} \text{ELBO}^{\text{AP}}(\boldsymbol{\Omega})$$

with objective function,

$$\begin{aligned}
\text{ELBO}^{\text{AP}}(\boldsymbol{\Omega}) &= \sum_{i=1}^{n}\sum_{j=1}^{p-1}\sum_{t=1}^{k} x_{ij}\left(\sqrt{\frac{2}{\pi}}\frac{\omega_{it}\alpha_{it}^{(s-1)}r_{jt}^{(s)}}{\sqrt{1+(\alpha_{it}^2)^{(s-1)}}}\right) \\
&\quad - \sum_{i=1}^{n} M_i \log\left(1 + \sum_{\ell=1}^{p-1}\left(1-\pi_{i\ell}^{(s)}\right)\exp\left(a_{0\ell}^{(s-1)} + \frac{1}{2}\left(C_{0\ell}^2\right)^{(s-1)} + \boldsymbol{L}_{i\ell}^{(s-1)}\right)\right) \\
&\quad - \sum_{i=1}^{n}\sum_{t=1}^{k}\left(\frac{\omega_{it}^2}{2} + \sqrt{\frac{2}{\pi}}\frac{\omega_{it}\xi_{it}^{(s)}\alpha_{it}^{(s-1)}}{\sqrt{1+(\alpha_{it}^2)^{(s-1)}}} - \sqrt{\frac{2}{\pi}}\frac{\alpha^*\omega_{it}\alpha_{it}^{(s-1)}}{\sqrt{1+(\alpha_{it}^2)^{(s-1)}}}\right) \\
&\quad + \sum_{i=1}^{n}\sum_{t=1}^{k}\left(\Psi_1^{AP}\left(\xi_{it}^{(s)}, \omega_{it}, \alpha_{it}^{(s-1)}\right) + \log\omega_{it}\right)
\end{aligned}$$

and gradient function,

$$\frac{\partial}{\partial \boldsymbol{\Omega}} \text{ELBO}^{\text{AP}}(\boldsymbol{\Omega}) = \sqrt{\frac{2}{\pi}}(X\boldsymbol{R}^{(s)}) \odot \left(\boldsymbol{A}^{(s-1)} \oslash \sqrt{1+(\boldsymbol{A}\odot\boldsymbol{A})^{(s-1)}}\right) - \widetilde{\boldsymbol{Q}}$$
$$- \left(\boldsymbol{\Omega} + \sqrt{\frac{2}{\pi}}\left(\boldsymbol{\Xi}^{(s)} \odot \boldsymbol{A}^{(s-1)}\right) \oslash \sqrt{1+(\boldsymbol{A}\odot\boldsymbol{A})^{(s-1)}} - (1 \oslash \boldsymbol{\Omega})\right)$$
$$+ \sqrt{\frac{2}{\pi}}\alpha^* \left(\boldsymbol{A}^{(s-1)} \oslash \sqrt{1+(\boldsymbol{A}\odot\boldsymbol{A})^{(s-1)}}\right) + \frac{\partial}{\partial \boldsymbol{\Omega}}\boldsymbol{\Psi}_1^{AP}\left(\boldsymbol{\Xi}^{(s)}, \boldsymbol{\Omega}, \boldsymbol{A}^{(s-1)}\right)$$

where, the elements of matrix $\widetilde{\boldsymbol{Q}}$,

$$\widetilde{Q}_{it} = M_i \left( \frac{\sum_{j=1}^{p-1}\left(1-\pi_{ij}^{(s)}\right) O_{ijt}^{(s-1)} \exp\left(a_{0j}^{(s-1)} + \frac{1}{2}\left(C_{0j}^2\right)^{(s-1)} + \boldsymbol{L}_{ij}^{(s-1)}\right)}{1+\sum_{\ell=1}^{p-1}\left(1-\pi_{i\ell}^{(s)}\right)\exp\left(a_{0\ell}^{(s-1)} + \frac{1}{2}\left(C_{0\ell}^2\right)^{(s-1)} + \boldsymbol{L}_{i\ell}^{(s-1)}\right)} \right)$$

and,

$$O_{ijt}^{(s-1)} = \frac{\partial}{\partial \omega_{it}} \boldsymbol{L}_{ij}^{(s-1)}$$

9. Update $\boldsymbol{A}^{(s)}$

$$\boldsymbol{A}^{(s)} = \arg\max_{\boldsymbol{A}} \text{ELBO}^{\text{AP}}(\boldsymbol{A})$$

with objective function,

$$\text{ELBO}^{\text{AP}}(\boldsymbol{A}) = \sum_{i=1}^{n}\sum_{j=1}^{p-1}\sum_{t=1}^{k} x_{ij}\left(\sqrt{\frac{2}{\pi}}\frac{\alpha_{it}\omega_{it}^{(s)} r_{jt}^{(s)}}{\sqrt{1+\alpha_{it}^2}}\right)$$
$$- \sum_{i=1}^{n} M_i \log\left(1+\sum_{\ell=1}^{p-1}\left(1-\pi_{i\ell}^{(s)}\right)\exp\left(a_{0\ell}^{(s-1)} + \frac{1}{2}\left(C_{0\ell}^2\right)^{(s-1)} + \boldsymbol{L}_{i\ell}^{(s-1)}\right)\right)$$
$$- \sum_{i=1}^{n}\sum_{t=1}^{k}\left(\sqrt{\frac{2}{\pi}}\frac{\alpha_{it}\omega_{it}^{(s)}\xi_{it}^{(s)}}{\sqrt{1+\alpha_{it}^2}} - \sqrt{\frac{2}{\pi}}\frac{\alpha^*\alpha_{it}\omega_{it}^{(s)}}{\sqrt{1+\alpha_{it}^2}} - \Psi_1^{AP}\left(\xi_{it}^{(s)}, \omega_{it}^{(s)}, \alpha_{it}\right) + \Psi_2^{AP}(\alpha_{it})\right)$$

and gradient function,

$$\frac{\partial}{\partial \boldsymbol{A}} \text{ELBO}^{\text{AP}}(\boldsymbol{A}) = \sqrt{\frac{2}{\pi}}\left(\left(\boldsymbol{\Omega}^{(s)} \odot \left(X\boldsymbol{R}^{(s)}\right)\right) \oslash (1+\boldsymbol{A}\odot\boldsymbol{A})^{\frac{3}{2}}\right) - \widetilde{\boldsymbol{Z}}$$
$$- \sqrt{\frac{2}{\pi}}\left(\left(\boldsymbol{\Omega}^{(s)} \odot \boldsymbol{\Xi}^{(s)}\right) \oslash (1+\boldsymbol{A}\odot\boldsymbol{A})^{\frac{3}{2}}\right)$$
$$+ \sqrt{\frac{2}{\pi}}\alpha^*\left(\boldsymbol{\Omega}^{(s)} \oslash (1+\boldsymbol{A}\odot\boldsymbol{A})^{\frac{3}{2}}\right)$$
$$+ \frac{\partial}{\partial \boldsymbol{A}}\boldsymbol{\Psi}_1^{AP}\left(\boldsymbol{\Xi}^{(s)}, \boldsymbol{\Omega}^{(s)}, \boldsymbol{A}\right) - \frac{\partial}{\partial \boldsymbol{A}}\boldsymbol{\Psi}_2^{AP}(\boldsymbol{A})$$

where, the elements of matrix, $\widetilde{\boldsymbol{Z}}$,

$$\widetilde{Z}_{it} = M_i \left( \frac{\sum_{j=1}^{p-1}\left(1-\pi_{ij}^{(s)}\right) A_{ijt}^{(s-1)} \exp\left(a_{0j}^{(s-1)} + \frac{1}{2}\left(C_{0j}^2\right)^{(s-1)} + \boldsymbol{L}_{ij}^{(s-1)}\right)}{1+\sum_{\ell=1}^{p-1}\left(1-\pi_{i\ell}^{(s)}\right)\exp\left(a_{0\ell}^{(s-1)} + \frac{1}{2}\left(C_{0\ell}^2\right)^{(s-1)} + \boldsymbol{L}_{i\ell}^{(s-1)}\right)} \right)$$

and,

$$A_{ijt}^{(s-1)} = \frac{\partial}{\partial \alpha_{it}} \boldsymbol{L}_{ij}^{(s-1)}$$

10. Update $\boldsymbol{A}_0^{(s)}$

$$\boldsymbol{A}_0^{(s)} = \arg\max_{\boldsymbol{A}_0} \text{ELBO}^{\text{AP}}(\boldsymbol{A}_0)$$

with objective function,

$$\text{ELBO}^{\text{AP}}(\boldsymbol{A}_0) = \sum_{i=1}^{n}\sum_{j=1}^{p-1} x_{ij}a_{0j} - \sum_{i=1}^{n} M_i \log\left(1 + \sum_{\ell=1}^{p-1}\left(1-\pi_{i\ell}^{(s)}\right)\exp\left(a_{0\ell} + \frac{1}{2}\left(C_{0\ell}^2\right)^{(s-1)} + \boldsymbol{L}_{i\ell}^{(s)}\right)\right)$$
$$- \frac{1}{2}\sum_{j=1}^{p-1} a_{0j}^2$$

and gradient function,

$$\frac{\partial}{\partial \boldsymbol{A}_0}\text{ELBO}^{\text{AP}}(\boldsymbol{A}_0) = X^T\mathbf{1}_n - \widetilde{\boldsymbol{C}} - \boldsymbol{A}_0$$

where $\mathbf{1}_n$ is an n-dimensional column vector of 1's and the elements of vector $\widetilde{\boldsymbol{C}}$,

$$\widetilde{C}_j = \sum_{i=1}^{n} M_i \left(\frac{\left(1-\pi_{ij}^{(s)}\right)\exp\left(a_{0j} + \frac{1}{2}\left(C_{0j}^2\right)^{(s-1)} + \boldsymbol{L}_{ij}^{(s)}\right)}{1 + \sum_{\ell=1}^{p-1}\left(1-\pi_{i\ell}^{(s)}\right)\exp\left(a_{0\ell} + \frac{1}{2}\left(C_{0\ell}^2\right)^{(s-1)} + \boldsymbol{L}_{i\ell}^{(s)}\right)}\right)$$

11. Update $\boldsymbol{C}_0^{(s)}$

$$\boldsymbol{C}_0^{(s)} = \arg\max_{\boldsymbol{C}_0} \text{ELBO}^{\text{AP}}(\boldsymbol{C}_0)$$

with objective function,

$$\text{ELBO}^{\text{AP}}(\boldsymbol{C}_0) = -\sum_{i=1}^{n} M_i \log\left(1 + \sum_{\ell=1}^{p-1}\left(1-\pi_{i\ell}^{(s)}\right)\exp\left(a_{0\ell}^{(s)} + \frac{1}{2}C_{0\ell}^2 + \boldsymbol{L}_{i\ell}^{(s)}\right)\right)$$
$$- \frac{1}{2}\left(\sum_{j=1}^{p-1} C_{0j}^2 - \log C_{0j}^2\right)$$

and gradient function,

$$\frac{\partial}{\partial \boldsymbol{C}_0}\text{ELBO}^{\text{AP}}(\boldsymbol{C}_0) = -\widetilde{\boldsymbol{D}} - \frac{1}{2}\left(\mathbf{1}_n - \frac{1}{\boldsymbol{C}_0}\right)$$

where, the elements of vector $\widetilde{\boldsymbol{D}}$,

$$\widetilde{D}_j = \sum_{i=1}^{n} M_i \left(\frac{\left(1-\pi_{ij}^{(s)}\right)\exp\left(a_{0j}^{(s)} + \frac{1}{2}C_{0j}^2 + \boldsymbol{L}_{ij}^{(s)}\right)}{1 + \sum_{\ell=1}^{p-1}\left(1-\pi_{i\ell}^{(s)}\right)\exp\left(a_{0\ell}^{(s)} + \frac{1}{2}C_{0\ell}^2 + \boldsymbol{L}_{i\ell}^{(s)}\right)}\right)$$

### 4 Derivatives of $\boldsymbol{L}_{ij}$ Function

Recall that,

$$\begin{aligned}
\boldsymbol{L_{ij}} &:= \sum_{t=1}^{k} \log L_{ijt} \\
&= \sum_{t=1}^{k} \log\left( \frac{2}{\sqrt{1-\lambda_{jt}^2\omega_{it}^2}} \times \exp\left\{ \frac{\lambda_{jt}^2\xi_{it}^2 + r_{jt}^2\omega_{it}^2 + 2r_{jt}\xi_{it}}{2(1-\lambda_{jt}^2\omega_{it}^2)} \right\} \times \varPhi\left[ \frac{(r_{jt}+\lambda_{jt}^2\xi_{it})\omega_{it}\alpha_{it}}{\sqrt{1-\lambda_{jt}^2\omega_{it}^2}\sqrt{1+\alpha_{it}^2-\lambda_{jt}^2\omega_{it}^2}} \right] \right) \\
&= \sum_{t=1}^{k} \left\{ \log 2 - \frac{1}{2}\log(1-\lambda_{jt}^2\omega_{it}^2) + \frac{\lambda_{jt}^2\xi_{it}^2 + r_{jt}^2\omega_{it}^2 + 2r_{jt}\xi_{it}}{2(1-\lambda_{jt}^2\omega_{it}^2)} + \log\left( \varPhi\left[ \frac{(r_{jt}+\lambda_{jt}^2\xi_{it})\omega_{it}\alpha_{it}}{\sqrt{1-\lambda_{jt}^2\omega_{it}^2}\sqrt{1+\alpha_{it}^2-\lambda_{jt}^2\omega_{it}^2}} \right] \right) \right\}
\end{aligned}$$

1. Differentiating $\boldsymbol{L_{ij}}$ with respect to $r_{jt}$,

$$\frac{\partial \boldsymbol{L_{ij}}}{\partial r_{jt}} = \frac{r_{jt}\omega_{it}^2 + \xi_{it}}{1-\lambda_{jt}^2\omega_{it}^2} + \frac{\phi\left( \frac{(r_{jt}+\lambda_{jt}^2\xi_{it})\omega_{it}\alpha_{it}}{\sqrt{1-\lambda_{jt}^2\omega_{it}^2}\sqrt{1+\alpha_{it}^2-\lambda_{jt}^2\omega_{it}^2}} \right)}{\varPhi\left[ \frac{(r_{jt}+\lambda_{jt}^2\xi_{it})\omega_{it}\alpha_{it}}{\sqrt{1-\lambda_{jt}^2\omega_{it}^2}\sqrt{1+\alpha_{it}^2-\lambda_{jt}^2\omega_{it}^2}} \right]} \times \frac{\omega_{it}\alpha_{it}}{\sqrt{1-\lambda_{jt}^2\omega_{it}^2}\sqrt{1+\alpha_{it}^2-\lambda_{jt}^2\omega_{it}^2}}$$

2. Differentiating $\boldsymbol{L_{ij}}$ with respect to $\lambda_{jt}^2$,

$$\begin{aligned}
\frac{\partial \boldsymbol{L_{ij}}}{\partial \lambda_{jt}^2} &= \frac{1}{2}\left( \frac{\omega_{it}^2}{1-\lambda_{jt}^2\omega_{it}^2} \right) + \frac{1}{2}\left[ \frac{\xi_{it}^2 + r_{jt}^2\omega_{it}^4 + 2r_{jt}\xi_{it}\omega_{it}^2}{(1-\lambda_{jt}^2\omega_{it}^2)^2} \right] + \frac{\phi\left( \frac{(r_{jt}+\lambda_{jt}^2\xi_{it})\omega_{it}\alpha_{it}}{\sqrt{1-\lambda_{jt}^2\omega_{it}^2}\sqrt{1+\alpha_{it}^2-\lambda_{jt}^2\omega_{it}^2}} \right)}{\varPhi\left[ \frac{(r_{jt}+\lambda_{jt}^2\xi_{it})\omega_{it}\alpha_{it}}{\sqrt{1-\lambda_{jt}^2\omega_{it}^2}\sqrt{1+\alpha_{it}^2-\lambda_{jt}^2\omega_{it}^2}} \right]} \\
&\times \left[ \frac{\xi_{it}\omega_{it}\alpha_{it}}{\sqrt{1-\lambda_{jt}^2\omega_{it}^2}\sqrt{1+\alpha_{it}^2-\lambda_{jt}^2\omega_{it}^2}} - \frac{(r_{jt}+\lambda_{jt}^2\xi_{it})\omega_{it}\alpha_{it}(-2\omega_{it}^2 - \omega_{it}^2\alpha_{it}^2 + 2\lambda_{jt}^2\omega_{it}^4)}{2[(1-\lambda_{jt}^2\omega_{it}^2)(1+\alpha_{it}^2-\lambda_{jt}^2\omega_{it}^2)]^{\frac{3}{2}}} \right]
\end{aligned}$$

3. Differentiating $\boldsymbol{L_{ij}}$ with respect to $\xi_{it}$,

$$\frac{\partial \boldsymbol{L_{ij}}}{\partial \xi_{it}} = \frac{\lambda_{jt}^2\xi_{it} + r_{jt}}{(1-\lambda_{jt}^2\omega_{it}^2)} + \frac{\phi\left( \frac{(r_{jt}+\lambda_{jt}^2\xi_{it})\omega_{it}\alpha_{it}}{\sqrt{1-\lambda_{jt}^2\omega_{it}^2}\sqrt{1+\alpha_{it}^2-\lambda_{jt}^2\omega_{it}^2}} \right)}{\varPhi\left[ \frac{(r_{jt}+\lambda_{jt}^2\xi_{it})\omega_{it}\alpha_{it}}{\sqrt{1-\lambda_{jt}^2\omega_{it}^2}\sqrt{1+\alpha_{it}^2-\lambda_{jt}^2\omega_{it}^2}} \right]} \times \frac{\lambda_{jt}^2\omega_{it}\alpha_{it}}{\sqrt{1-\lambda_{jt}^2\omega_{it}^2}\sqrt{1+\alpha_{it}^2-\lambda_{jt}^2\omega_{it}^2}}$$

4. Differentiating $\boldsymbol{L_{ij}}$ with respect to $\omega_{it}$,

$$\begin{aligned}
\frac{\partial \boldsymbol{L_{ij}}}{\partial \omega_{it}} &= \frac{\lambda_{jt}^2\omega_{it}}{1-\lambda_{jt}^2\omega_{it}^2} + \frac{r_{jt}^2\omega_{it} + \lambda_{jt}^4\omega_{it}\xi_{it}^2 + 2\lambda_{jt}^2\omega_{it}r_{jt}\xi_{it}}{(1-\lambda_{jt}^2\omega_{it}^2)^2} + \frac{\phi\left( \frac{(r_{jt}+\lambda_{jt}^2\xi_{it})\omega_{it}\alpha_{it}}{\sqrt{1-\lambda_{jt}^2\omega_{it}^2}\sqrt{1+\alpha_{it}^2-\lambda_{jt}^2\omega_{it}^2}} \right)}{\varPhi\left[ \frac{(r_{jt}+\lambda_{jt}^2\xi_{it})\omega_{it}\alpha_{it}}{\sqrt{1-\lambda_{jt}^2\omega_{it}^2}\sqrt{1+\alpha_{it}^2-\lambda_{jt}^2\omega_{it}^2}} \right]} \\
&\times \left[ \frac{(r_{jt}+\lambda_{jt}^2\xi_{it})\alpha_{it}}{\sqrt{1-\lambda_{jt}^2\omega_{it}^2}\sqrt{1+\alpha_{it}^2-\lambda_{jt}^2\omega_{it}^2}} - \frac{(r_{jt}+\lambda_{jt}^2\xi_{it})\omega_{it}\alpha_{it}(-2\lambda_{jt}^2\omega_{it} - \lambda_{jt}^2\omega_{it}\alpha_{it}^2 + 2\lambda_{jt}^4\omega_{it}^3)}{[(1-\lambda_{jt}^2\omega_{it}^2)(1+\alpha_{it}^2-\lambda_{jt}^2\omega_{it}^2)]^{\frac{3}{2}}} \right]
\end{aligned}$$

5. Differentiating $\boldsymbol{L_{ij}}$ with respect to $\alpha_{it}$,

$$\frac{\partial \boldsymbol{L_{ij}}}{\partial \alpha_{it}} = \frac{\phi\left(\frac{(r_{jt}+\lambda_{jt}^2\xi_{it})\omega_{it}\alpha_{it}}{\sqrt{1-\lambda_{jt}^2\omega_{it}^2}\sqrt{1+\alpha_{it}^2-\lambda_{jt}^2\omega_{it}^2}}\right)}{\Phi\left[\frac{(r_{jt}+\lambda_{jt}^2\xi_{it})\omega_{it}\alpha_{it}}{\sqrt{1-\lambda_{jt}^2\omega_{it}^2}\sqrt{1+\alpha_{it}^2-\lambda_{jt}^2\omega_{it}^2}}\right]}$$
$$\times\left[\frac{(r_{jt}+\lambda_{jt}^2\xi_{it})\omega_{it}}{\sqrt{1-\lambda_{jt}^2\omega_{it}^2}\sqrt{1+\alpha_{it}^2-\lambda_{jt}^2\omega_{it}^2}} - \frac{(r_{jt}+\lambda_{jt}^2\xi_{it})\omega_{it}\alpha_{it}(\alpha_{it}-\lambda_{jt}^2\omega_{it}^2\alpha_{it})}{[(1-\lambda_{jt}^2\omega_{it}^2)(1+\alpha_{it}^2-\lambda_{jt}^2\omega_{it}^2)]^{\frac{3}{2}}}\right]$$

### 5 Taylor Expansion and it's gradients for $\Psi_1(\xi_{it}, \omega_{it}, \alpha_{it})$

Let $F_{it} \sim \mathcal{SN}(\xi_{it}, \omega_{it}, \alpha_{it})$. We need to find $\Psi_1(\xi_{it}, \omega_{it}, \alpha_{it}) = \mathbb{E}[\log \Phi(\alpha_0(F_{it} - \alpha^*))]$ where $\alpha^* = -\sqrt{\frac{2}{\pi}}\frac{\alpha_0}{\sqrt{1+\alpha_0^2}}$. Let,

$$\alpha_0(F_{it} - \alpha^*) = X_{it}$$

Then,

$$\mu_{it} = \mathbb{E}[X_{it}] = \alpha_0\left(\xi_{it} + \frac{\omega_{it}\alpha_{it}}{\sqrt{1+\alpha_{it}^2}}\sqrt{\frac{2}{\pi}} - \alpha^*\right)$$
$$\sigma_{it}^2 = Var(X_{it}) = \alpha_0^2\omega_{it}^2\left(1 - \frac{2}{\pi}\frac{\alpha_{it}^2}{1+\alpha_{it}^2}\right)$$

We now find the $\mathbb{E}[\log \Phi(X_{it})]$.

Since, there is no closed form available for the above, we approximate it using a Taylor expansion.

Let, $f(X_{it}) = \log \Phi(X_{it})$. Given $\mu_{it}$ and $\sigma_{it}^2$, a Taylor expansion of the expected value of $f(X_{it})$ is given by,

$$\begin{aligned}
\mathbb{E}(f(X_{it})) &= \mathbb{E}(f(\mu_{it} + (X_{it} - \mu_{it}))) \\
&\approx \mathbb{E}\left(f(\mu_{it}) + f'(\mu_{it})(X_{it} - \mu_{it}) + \frac{1}{2}f''(\mu_{it})(X_{it} - \mu_{it})^2\right) \\
&= f(\mu_{it}) + f'(\mu_{it})\mathbb{E}[X_{it} - \mu_{it}] + \frac{1}{2}f''(\mu_{it})\mathbb{E}\left[(X_{it} - \mu_{it})^2\right] \\
&= f(\mu_{it}) + \frac{f''(\mu_{it})}{2}\sigma_{it}^2
\end{aligned}$$

Therefore,

$$\mathbb{E}(\log \Phi[X_{it}]) \approx \Psi_1^{AP}(\xi_{it}, \omega_{it}, \alpha_{it})$$

where,

$$\begin{aligned}
\Psi_1^{AP}(\xi_{it},\omega_{it},\alpha_{it}) &= \log \Phi[\mu_{it}] + \frac{f''(\mu_{it})}{2}\sigma_{it}^2 \\
f''(\mu_{it}) &= -M_1 - M_2 \\
M_1 &= \frac{1}{\sqrt{2\pi}} \times \mu_{it} \times \exp\left(\frac{-\mu_{it}^2}{2}\right) \times \frac{1}{\Phi[\mu_{it}]} \\
M_2 &= \left(\frac{\phi(\mu_{it})}{\Phi[\mu_{it}]}\right)^2 = \frac{1}{2\pi}\frac{\exp\left(-\mu_{it}^2\right)}{\left(\Phi[\mu_{it}]\right)^2}
\end{aligned}$$

Now,

1. Differentiating $\Psi_1^{AP}(\xi_{it},\omega_{it},\alpha_{it})$ with respect to $\xi_{it}$,

$$\frac{\partial}{\partial \xi_{it}}\Psi_1^{AP}(\xi_{it},\omega_{it},\alpha_{it}) = \frac{\phi(\mu_{it})}{\Phi[\mu_{it}]} \times \frac{\partial \mu_{it}}{\partial \xi_{it}} + \frac{\alpha_0^2\omega_{it}^2}{2}\left(1 - \frac{2}{\pi}\frac{\alpha_{it}^2}{1+\alpha_{it}^2}\right)\left(-\frac{\partial M_1}{\partial \xi_{it}} - \frac{\partial M_2}{\partial \xi_{it}}\right)$$

2. Differentiating $\Psi_1^{AP}(\xi_{it},\omega_{it},\alpha_{it})$ with respect to $\omega_{it}$,

$$\begin{aligned}
\frac{\partial}{\partial \omega_{it}}\Psi_1^{AP}(\xi_{it},\omega_{it},\alpha_{it}) = {}& \frac{\phi(\mu_{it})}{\Phi[\mu_{it}]} \times \frac{\partial \mu_{it}}{\partial \omega_{it}} + \alpha_0^2\omega_{it}\left(1 - \frac{2}{\pi}\frac{\alpha_{it}^2}{1+\alpha_{it}^2}\right)(-M_1 - M_2) \\
&+ \frac{\alpha_0^2\omega_{it}^2}{2}\left(1 - \frac{2}{\pi}\frac{\alpha_{it}^2}{1+\alpha_{it}^2}\right)\left(-\frac{\partial M_1}{\partial \omega_{it}} - \frac{\partial M_2}{\partial \omega_{it}}\right)
\end{aligned}$$

3. Differentiating $\Psi_1^{AP}(\xi_{it},\omega_{it},\alpha_{it})$ with respect to $\alpha_{it}$,

$$\begin{aligned}
\frac{\partial}{\partial \alpha_{it}}\Psi_1^{AP}(\xi_{it},\omega_{it},\alpha_{it}) = {}& \frac{\phi(\mu_{it})}{\Phi[\mu_{it}]} \times \frac{\partial \mu_{it}}{\partial \alpha_{it}} + \frac{\alpha_0^2\omega_{it}^2}{2}\left(-\frac{\partial}{\partial \alpha_{it}}\left(\frac{2}{\pi}\frac{\alpha_{it}^2}{1+\alpha_{it}^2}\right)\right)(-M_1 - M_2) \\
&+ \frac{\alpha_0^2\omega_{it}^2}{2}\left(1 - \frac{2}{\pi}\frac{\alpha_{it}^2}{1+\alpha_{it}^2}\right)\left(-\frac{\partial M_1}{\partial \alpha_{it}} - \frac{\partial M_2}{\partial \alpha_{it}}\right)
\end{aligned}$$

### 6 Taylor Expansion and it's gradients for $\Psi_2(\alpha_{it})$

Let $\mu_{it}^{(V)}$ and $\sigma_{it}^{2(V)}$ be the mean and variance of $V_{it} = \alpha_{it}U_{it}$, respectively, where $U_{it} \sim \mathcal{SN}(0, 1, \alpha_{it})$, i.e.,

$$\mu_{it}^{(V)} = \frac{\alpha_{it}^2}{\sqrt{1+\alpha_{it}^2}}\sqrt{\frac{2}{\pi}}$$

$$\sigma_{it}^{2(V)} = \alpha_{it}^2 - \left(\mu_{it}^{(V)}\right)^2$$

We need to find $\Psi_2(\alpha_{it}) = \mathbb{E}(\log \Phi[V_{it}])$.

Since, there is no closed form available for the above integral, we approximate it using a Taylor expansion.

$$\mathbb{E}[\log \Phi(V_{it})] \approx \Psi_2^{AP}(\alpha_{it})$$

where,

$$\Psi_2^{AP}(\alpha_{it}) = \log \Phi\left[\mu_{it}^{(V)}\right] + \frac{1}{2}\left(\alpha_{it}^2 - \left(\mu_{it}^{(V)}\right)^2\right)(-N_1 - N_2)$$

$$N_1 = \frac{1}{\sqrt{2\pi}} \times \mu_{it}^{(V)} \times \exp\left(-\frac{1}{2}\left(\mu_{it}^{(V)}\right)^2\right) \times \frac{1}{\Phi\left[\mu_{it}^{(V)}\right]}$$

$$N_2 = \frac{1}{2\pi}\frac{\exp\left(-\left(\mu_{it}^{(V)}\right)^2\right)}{\left(\Phi\left[\mu_{it}^{(V)}\right]\right)^2}$$

Now, differentiating $\Psi_2^{AP}(\alpha_{it})$ with respect to $\alpha_{it}$,

$$\frac{\partial}{\partial \alpha_{it}}\Psi_2^{AP}(\alpha_{it}) = \frac{\phi\left(\mu_{it}^{(V)}\right)}{\Phi\left[\mu_{it}^{(V)}\right]} \times \frac{\partial \mu_{it}^{(V)}}{\partial \alpha_{it}} + \left(\alpha_{it} - \mu_{it}^{(V)}\frac{\partial \mu_{it}^{(V)}}{\partial \alpha_{it}}\right)(-N_1 - N_2)$$
$$+ \frac{1}{2}\left(\alpha_{it}^2 - \left(\mu_{it}^{(V)}\right)^2\right)\left(-\frac{\partial N_1}{\partial \alpha_{it}} - \frac{\partial N_2}{\partial \alpha_{it}}\right)$$

#### 7 Update for $\pi_{ij}$ parameter

Consider a Zero Inflated Factor Analysis Logistic Skew Normal Poisson (ZIFA - LSNP) model having,

$$\text{observation space, } x_{ij} \mid \mu_{ij}, z_{ij} \overset{\text{ind}}{\sim} \begin{cases} \text{Poisson}(\mu_{ij}) & \text{if } z_{ij} = 0 \\ 0 & \text{if } z_{ij} = 1 \end{cases}$$

$$\text{parameter space, } \log \mu_{ij} = \varUpsilon_{i0} + \beta_{0j} + \boldsymbol{F}_i^T \boldsymbol{\beta_j}$$

with priors,

$$z_{ij} \overset{\text{ind}}{\sim} \mathcal{B}\text{ern}(\kappa_j), \qquad \kappa_j \overset{\text{iid}}{\sim} \mathcal{B}\text{eta}(\nu_1, \nu_2),$$

$$\beta_{0j} \overset{\text{iid}}{\sim} \mathcal{N}(0, 1), \qquad F_{it} \overset{\text{iid}}{\sim} \mathcal{SN}(\alpha^*, 1, \alpha_0),$$

$$\beta_{jt} \overset{\text{ind}}{\sim} \mathcal{N}(0, \delta_{jt}^{-1}), \qquad \delta_{jt} \overset{\text{iid}}{\sim} \mathcal{G}\text{amma}(g_1, g_2).$$

Now,

$$\begin{aligned} p(\boldsymbol{x_i} \mid \mu_{ij}, z_{ij}) &= \prod_{j=1}^{p} p(x_{ij} \mid \mu_{ij}, z_{ij}) \\ &= \prod_{j=1}^{p} \left[ (\text{Poisson}(\mu_{ij}))^{1-z_{ij}} (\mathbb{I}(x_{ij} = 0))^{z_{ij}} \right] \\ \log p(\boldsymbol{x_i} \mid \mu_{ij}, z_{ij}) &= \sum_{j=1}^{p} (1 - z_{ij}) \left[ x_{ij} \log \mu_{ij} - \mu_{ij} - \log x_{ij}! \right] \\ &= \sum_{j=1}^{p} (1 - z_{ij}) \left[ x_{ij} \left( \varUpsilon_{i0} + \beta_{0j} + \sum_{t=1}^{k} F_{it} \beta_{jt} \right) - \exp\left( \varUpsilon_{i0} + \beta_{0j} + \boldsymbol{F}_i^T \boldsymbol{\beta}_j \right) - \log x_{ij}! \right] \end{aligned}$$

Hence,

$$
\begin{aligned}
\mathrm{ELBO}^{\mathrm{POI}} &= \sum_{i=1}^{n}\sum_{j=1}^{p}(1-\pi_{ij})\left[x_{ij}\left(\varUpsilon_{i0}+a_{0j}+\sum_{t=1}^{k}\left(\xi_{it}+\frac{\omega_{it}\alpha_{it}}{\sqrt{1+\alpha_{it}^2}}\sqrt{\frac{2}{\pi}}\right)(r_{jt})\right)\right] \\
&-\sum_{i=1}^{n}\sum_{j=1}^{p}(1-\pi_{ij})\left[\exp\left(\varUpsilon_{i0}+a_{0j}+\frac{1}{2}C_{0j}^2+\boldsymbol{L}_{ij}\right)+\log x_{ij}!\right] \\
&+\sum_{j=1}^{p}\sum_{t=1}^{k}\left(\left(g_1-\frac{1}{2}\right)(\psi(g_{1jt})-\log g_{2jt})-g_2\frac{g_{1jt}}{g_{2jt}}-\frac{1}{2}\left(\lambda_{jt}^2+r_{jt}^2\right)\frac{g_{1jt}}{g_{2jt}}\right) \\
&+\sum_{j=1}^{p}\sum_{t=1}^{k}\left(\frac{1}{2}\log\lambda_{jt}^2+g_{1jt}-\log g_{2jt}+\log\varGamma g_{1jt}+(1-g_{1jt})\psi(g_{1jt})\right) \\
&+\sum_{j=1}^{p}[(\nu_1-\tau_{j1})\{\psi(\tau_{j1})-\psi(\tau_{j1}+\tau_{j2})\}+(\nu_2-\tau_{j2})\{\psi(\tau_{j2})-\psi(\tau_{j1}+\tau_{j2})\} \\
&+\log\mathcal{B}(\tau_{j1},\tau_{j2})]+\sum_{i=1}^{n}\sum_{j=1}^{p}[\pi_{ij}\{\psi(\tau_{j1})-\psi(\tau_{j1}+\tau_{j2})-\log\pi_{ij}\} \\
&+(1-\pi_{ij})\{\psi(\tau_{j2})-\psi(\tau_{j1}+\tau_{j2})-\log(1-\pi_{ij})\}]-\sum_{i=1}^{n}\sum_{t=1}^{k}\left[\frac{\omega_{it}^2}{2}+\frac{\xi_{it}^2}{2}+\sqrt{\frac{2}{\pi}}\frac{\xi_{it}\omega_{it}\alpha_{it}}{\sqrt{1+\alpha_{it}^2}}\right. \\
&\left.-\alpha^*\left(\xi_{it}+\frac{\omega_{it}\alpha_{it}}{\sqrt{1+\alpha_{it}^2}}\sqrt{\frac{2}{\pi}}\right)-\varPsi(\xi_{it},\omega_{it},\alpha_{it})-\log\omega_{it}+\varPsi(\alpha_{it})\right]
\end{aligned}
$$

Taking the first and second partial derivative of $\mathrm{ELBO}^{\mathrm{POI}}$ with respect to $\varUpsilon_{i0}$, we get,

$$
\frac{\partial}{\partial\varUpsilon_{i0}}\mathrm{ELBO}^{\mathrm{POI}}=\sum_{j=1}^{p}(1-\pi_{ij})\left[x_{ij}-\exp\left(\varUpsilon_{i0}+a_{0j}+\frac{1}{2}C_{0j}^2+\boldsymbol{L}_{ij}\right)\right]
$$

and,

$$
\frac{\partial^2}{\partial\varUpsilon_{i0}^2}\mathrm{ELBO}^{\mathrm{POI}}=-\sum_{j=1}^{p}(1-\pi_{ij})\exp\left(\varUpsilon_{i0}+a_{0j}+\frac{1}{2}C_{0j}^2+\boldsymbol{L}_{ij}\right)<0
$$

Thus, $\mathrm{ELBO}^{\mathrm{POI}}$ has a unique maximizer,

$$
\hat{\varUpsilon}_{i0}=\log\left\{\frac{M_i}{\sum_{j=1}^{p}(1-\pi_{ij})\exp\left(a_{0j}+\frac{1}{2}C_{0j}^2+\boldsymbol{L}_{ij}\right)}\right\}
$$

Plugging in $\hat{\Upsilon}_{i0}$ for $\Upsilon_{i0}$ yields,

$$
\begin{aligned}
\text{ELBO}^{\text{POI}}_{\hat{\Upsilon}_i} &= \sum_{i=1}^{n}\sum_{j=1}^{p} x_{ij}a_{0j} + \sum_{i=1}^{n}\sum_{j=1}^{p}\sum_{t=1}^{k} x_{ij}r_{jt}\left(\xi_{it} + \frac{\omega_{it}\alpha_{it}}{\sqrt{1+\alpha_{it}^2}}\sqrt{\frac{2}{\pi}}\right) \\
&- \sum_{i=1}^{n} M_i \log\left(\sum_{j=1}^{p}(1-\pi_{ij})\exp\left(a_{0j} + \frac{1}{2}C_{0j}^2 + \boldsymbol{L}_{ij}\right)\right) \\
&+ \sum_{j=1}^{p}\sum_{t=1}^{k}\left(\left(g_1 - \frac{1}{2}\right)(\psi(g_{1jt}) - \log g_{2jt}) - g_2\frac{g_{1jt}}{g_{2jt}} - \frac{1}{2}\left(\lambda_{jt}^2 + r_{jt}^2\right)\frac{g_{1jt}}{g_{2jt}}\right) \\
&+ \sum_{j=1}^{p}\sum_{t=1}^{k}\left(\frac{1}{2}\log\lambda_{jt}^2 + g_{1jt} - \log g_{2jt} + \log \Gamma g_{1jt} + (1-g_{1jt})\psi(g_{1jt})\right) \\
&+ \sum_{j=1}^{p}\Bigg[(\nu_1 - \tau_{j1})\{\psi(\tau_{j1}) - \psi(\tau_{j1}+\tau_{j2})\} + (\nu_2 - \tau_{j2})\{\psi(\tau_{j2}) - \psi(\tau_{j1}+\tau_{j2})\} \\
&+ \log\mathcal{B}(\tau_{j1},\tau_{j2})\Bigg] + \sum_{i=1}^{n}\sum_{j=1}^{p}\Bigg[\pi_{ij}\{\psi(\tau_{j1}) - \psi(\tau_{j1}+\tau_{j2}) - \log\pi_{ij}\} \\
&+ (1-\pi_{ij})\{\psi(\tau_{j2}) - \psi(\tau_{j1}+\tau_{j2}) - \log(1-\pi_{ij})\}\Bigg] \\
&- \sum_{i=1}^{n}\sum_{t=1}^{k}\Bigg[\frac{\omega_{it}^2}{2} + \frac{\xi_{it}^2}{2} + \sqrt{\frac{2}{\pi}}\frac{\xi_{it}\omega_{it}\alpha_{it}}{\sqrt{1+\alpha_{it}^2}} - \alpha^*\left(\xi_{it} + \frac{\omega_{it}\alpha_{it}}{\sqrt{1+\alpha_{it}^2}}\sqrt{\frac{2}{\pi}}\right) \\
&- \Psi_1(\xi_{it},\omega_{it},\alpha_{it}) - \log\omega_{it} + \Psi_2(\alpha_{it})\Bigg].
\end{aligned}
$$

Now, to find an update for $\pi_{ij}$ we take the first and second partial derivative of $\text{ELBO}^{\text{POI}}$ with respect to $\pi_{ij}$,

$$
\begin{aligned}
\frac{\partial}{\partial\pi_{ij}}\text{ELBO}^{\text{POI}} &= \psi(\tau_{j1}) - \psi(\tau_{j2}) + \log\left(\frac{1-\pi_{ij}}{\pi_{ij}}\right) + \exp\left(a_{0j} + \frac{1}{2}C_{0j}^2 + \boldsymbol{L}_{ij}\right) \\
&- x_{ij}\left(\Upsilon_{i0} + a_{0j} + \sum_{t=1}^{k}\left(\xi_{it} + \frac{\omega_{it}\alpha_{it}}{\sqrt{1+\alpha_{it}^2}}\sqrt{\frac{2}{\pi}}\right)(r_{jt})\right)
\end{aligned}
$$

and,

$$
\frac{\partial^2}{\partial\pi_{ij}^2}\text{ELBO}^{\text{POI}} = -\frac{1}{\pi_{ij}(1-\pi_{ij})} < 0
$$

Thus,

$$
\hat{\pi}_{ij} = \begin{cases} \frac{\exp(\psi(\tau_{j1}))}{\exp(\psi(\tau_{j1})) + \exp\left(\psi(\tau_{j2}) - \exp\left(\Upsilon_{i0} + a_{0j} + \frac{1}{2}C_{0j}^2 + \boldsymbol{L}_{ij}\right)\right)} & \text{if } x_{ij} = 0 \\ 0 & \text{if } x_{ij} > 0 \end{cases}
$$

### 8 Skewness Plots

#### 8.1 Simulation plots

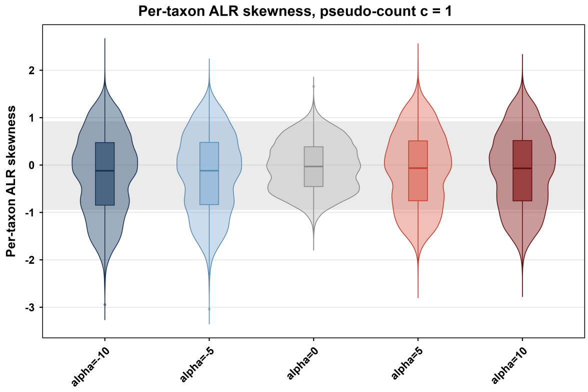


(a) Per-Taxon ALR-transformed Skewness after adding 1 as pseudo-count

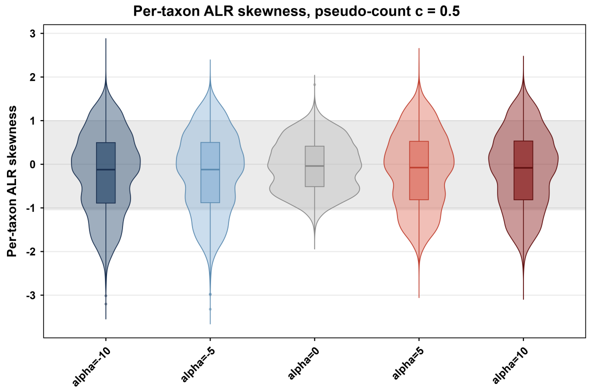


(b) Per-Taxon ALR-transformed Skewness after adding 0.5 as pseudo-count

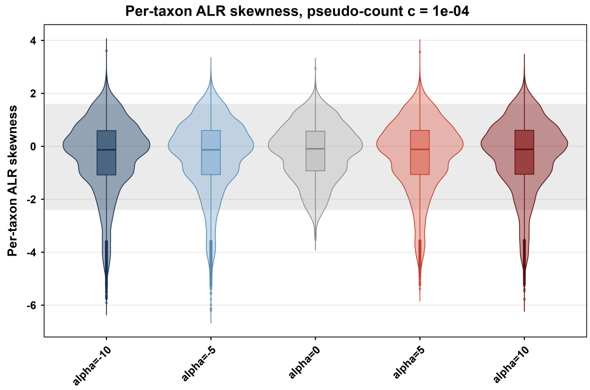


(c) Per-Taxon ALR-transformed Skewness after adding $10^{-4}$ as pseudo-count

Figure 1

#### 8.2 Real data plots

In this section, we compute the sample skewness of the ALR-transformed counts for each taxon in the real data. Because the ALR transformation is undefined at zero, a small pseudo-count $c \in \{1, 0.5, 10^{-2}, 10^{-4}\}$ was added to all counts before transformation. This ad hoc adjustment is used only for this descriptive summary. The ZIFA-LSNM model does not need it, because it handles zeros explicitly through its zero-inflation component. As shown in Figures 2(a)–2(d), roughly half of the genera have skewed ALR-transformed counts: 52.6%, 53.4%, 45.8% and 45.0% of taxa have an absolute sample skewness greater than 0.5 for $c = 1, 0.5, 10^{-2}$ and $10^{-4}$, respectively. Sample skewness was computed with the `skewness()` function of the `moments` package in `R`.

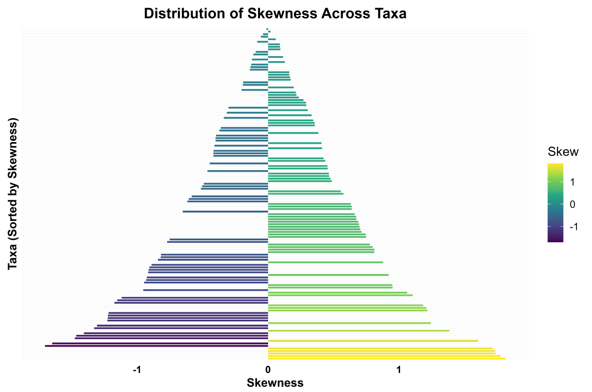


(a) Per-Taxon ALR-transformed Skewness after adding 1 as pseudo-count

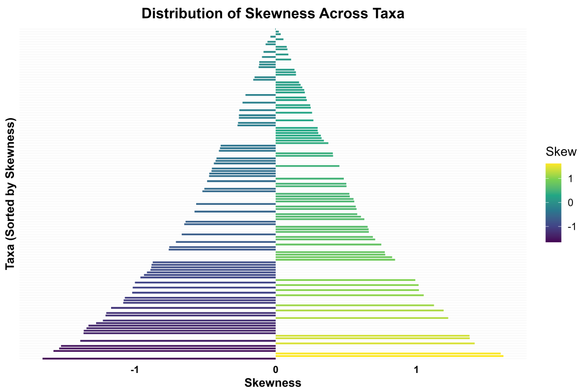


(b) Per-Taxon ALR-transformed Skewness after adding 0.5 as pseudo-count

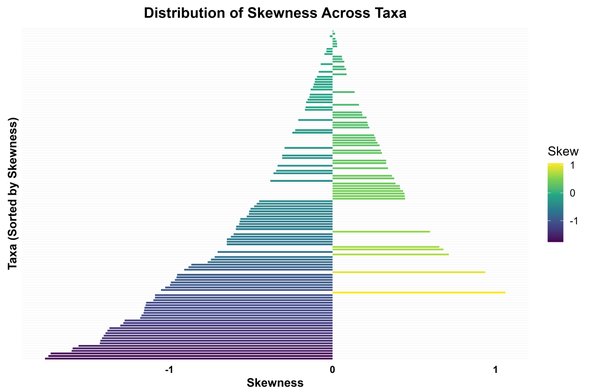


(c) Per-Taxon ALR-transformed Skewness after adding $10^{-2}$ as pseudo-count

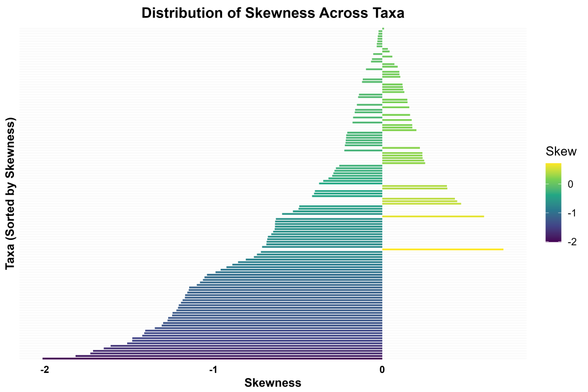


(d) Per-Taxon ALR-transformed Skewness after adding $10^{-4}$ as pseudo-count

Figure 2
